# Interpreting Neural Networks for Biological Sequences by Learning Stochastic Masks

**DOI:** 10.1101/2021.04.29.441979

**Authors:** Johannes Linder, Alyssa La Fleur, Zibo Chen, Ajasja Ljubetič, David Baker, Sreeram Kannan, Georg Seelig

## Abstract

Sequence-based neural networks can learn to make accurate predictions from large biological datasets, but model interpretation remains challenging. Many existing feature attribution methods are optimized for continuous rather than discrete input patterns and assess individual feature importance in isolation, making them ill-suited for interpreting non-linear interactions in molecular sequences. Building on work in computer vision and natural language processing, we developed an approach based on deep generative modeling - Scrambler networks - wherein the most salient sequence positions are identified with learned input masks. Scramblers learn to generate Position-Specific Scoring Matrices (*PSSMs*) where unimportant nucleotides or residues are ‘scrambled’ by raising their entropy. We apply Scramblers to interpret the effects of genetic variants, uncover non-linear interactions between cis-regulatory elements, explain binding specificity for protein-protein interactions, and identify structural determinants of *de novo* designed proteins. We show that interpretation based on a generative model allows for efficient attribution across large datasets and results in high-quality explanations, often outperforming state-of-the-art methods.

## Introduction

Deep Neural Networks have successfully been applied to a diverse set of biological sequence prediction problems, including predicting transcription factor binding (Alipanahi et al., 2015; Avsec et al., 2021; Eraslan et al., 2019; Movva et al., 2019), chromatin modification and accessibility (Zhou et al., 2015), RNA processing (Arefeen et al., 2019; Bogard et al., 2019; Cheng et al., 2019; Jaganathan et al., 2019), translation regulation (Sample et al., 2019) and protein structure (Senior et al., 2020, Yang et al., 2020). Neural networks excel at learning complex relationships from large datasets without requiring much tuning, but interpreting their predictions is challenging because the learned regulatory logic is embedded deep within the network layers. Nevertheless, interpretability is necessary if we want to connect network predictions to established biology, learn new regulatory rules or validate sequence designs (Talukder et al., 2020; Lanchantin et al., 2016; Schreiber et al., 2020; Norn et al., 2020; Kelley et al., 2016; Zeng et al., 2018; Kelley et al., 2019; Zeng et al., 2020; Singh et al., 2019; Avsec et al., 2021).

Nonlinear interactions are abundant between biological sequence features. Such dependencies make it necessary to consider groups of features and their surrounding context when interpreting a model prediction instead of independently evaluating each feature in isolation. However, many current neural network attribution methods rely on local perturbations, limiting their ability to identify such relationships. For example, 5’ UTRs can contain multiple upstream start codons (uAUGs; Calvo et al., 2009; Araujo et al., 2012; Whiffin et al., 2020) preceding the primary start. Which AUG is chosen as the start of translation depends on the extent to which the surrounding context – e.g., competing AUGs – represses its usage. Two nearby uAUGs may even “hide” each other, as each AUG is capable of repressing translation initiation at the primary start independently. In-silico saturation mutagenesis – which systematically exchanges one nucleotide and approximates its importance by predicted functional change – would incorrectly conclude both uAUGs are irrelevant: Neither uAUG would be identified as repressive, as knocking down only one would not change the prediction. Other local approximation methods face similar problems, such as those basing their estimation on gradients (Simonyan et al., 2013; Zeiler et al., 2014; Springenberg et al., 2014; Sundararajan et al., 2017; Shrikumar et al., 2017; Lundberg et al., 2017) or local linear models (Ribeiro et al., 2016).

In this paper, we introduce a feature attribution approach tailored for biological sequence predictors and based on previous work in learning interpretable input masks (Fong et al., 2017, Fong et al., 2019; Dabkowski et al., 2017; Zintgraf et al., 2017; Chen et al., 2018; Yoon et al., 2018; Chang et al., 2018; Carter et al., 2019; Covert et al., 2020). We train a deep generative model, referred to as the Scrambler Network, to learn masks that include only the smallest set of features in an input sequence necessary to reconstruct the original prediction. A mask that fulfills the inclusion objective overcomes the issues with local approximation mentioned above. For example, such a mask would learn to include one of the uAUGs from the 5’ UTR example to preserve the repressive prediction. In a complementary formulation, we optimize masks to occlude the smallest set of features which destroy the prediction. These features may overlap with those identified by inclusion masks but are not necessarily identical; for the 5’ UTR example, the occlusion mask would need to occlude both of the uAUGS to destroy the repressive prediction.

The earliest mask-based attribution methods masked inputs by either fading or blurring (Fong et al., 2017; Dabkowski et al., 2017), or they replaced features with zeros (Chen et al., 2018; Yoon et al., 2018), which is ill-suited for one-hot encoded sequence patterns. More recent methods replace masked input features with random samples or counterfactual values to keep masked inputs in the distribution of valid predictor patterns (Chang et al., 2018; Carter et al., 2019). However, these recent methods are based on individually optimizing each mask rather than learning a generative model. Here, we combine ideas from generative modeling with probabilistic masking to interpret biological sequences. Specifically, Scramblers learn to output Position-Specific Scoring Matrices *(PSSMs*) with minimal (inclusion) or maximal (occlusion) conservation such that discrete samples either reconstruct or destroy the prediction. A sequence position is ‘masked’ by raising the entropy, or *temperature*, of the underlying feature distribution such that samples sent to the predictor become less informative, and the Scrambler is trained by backpropagation using a gradient estimator. We show that Scramblers avoid overfitting to spurious per-example signals, which can otherwise be problematic for non-generative methods. Furthermore, using a generative model allows us to interpret new patterns more efficiently than per-example masking methods. See Table S1 for a summary comparison of masking methods.

In addition to introducing generative, probabilistic masks to biological sequences, we develop several improvements for mask-based interpretation. For example, Scramblers can discover multiple salient feature sets within a single input pattern using mask dropout- and bias layers, enabling users to specify features to include or exclude in the solution at inference time. In addition to finding reconstructive features, we alternatively optimize the Scrambler to find enhancing or repressive features that either maximize or minimize the prediction. We also explore different architectures for interpreting pairwise predictors in the context of protein-protein interaction. Finally, to reduce interpretation artifacts for predictors where input patterns contain a highly variable number of features, we apply per-example optimization as a fine-tuning step to the initial Scrambler outputs. Throughout the paper, we show that Scramblers learn meaningful feature attributions for several visual- and sequence-predictive tasks, including RNA regulation, protein-protein interaction and protein tertiary structure, often outperforming other interpretation methods.

## Results

### Scrambling Networks

The Scrambler architecture is illustrated in Figure 1A (left). Given a differentiable pre-trained predictor 𝒫 and a one-hot encoded input pattern ***x*** ∈ {0, 1}^*N ×M*^ (representing an *N* -length sequence), we let a trainable network 𝒮 called the *Scrambler* generate a set of real-valued importance scores 𝒮 (***x***) ∈ (0, ∞]^*N*^. These scores are used in Equation 1 below to produce a probability distribution which interpolates between the input pattern ***x*** and a non-informative background distribution 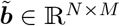(Figure 1B-C).

**Figure 1:**
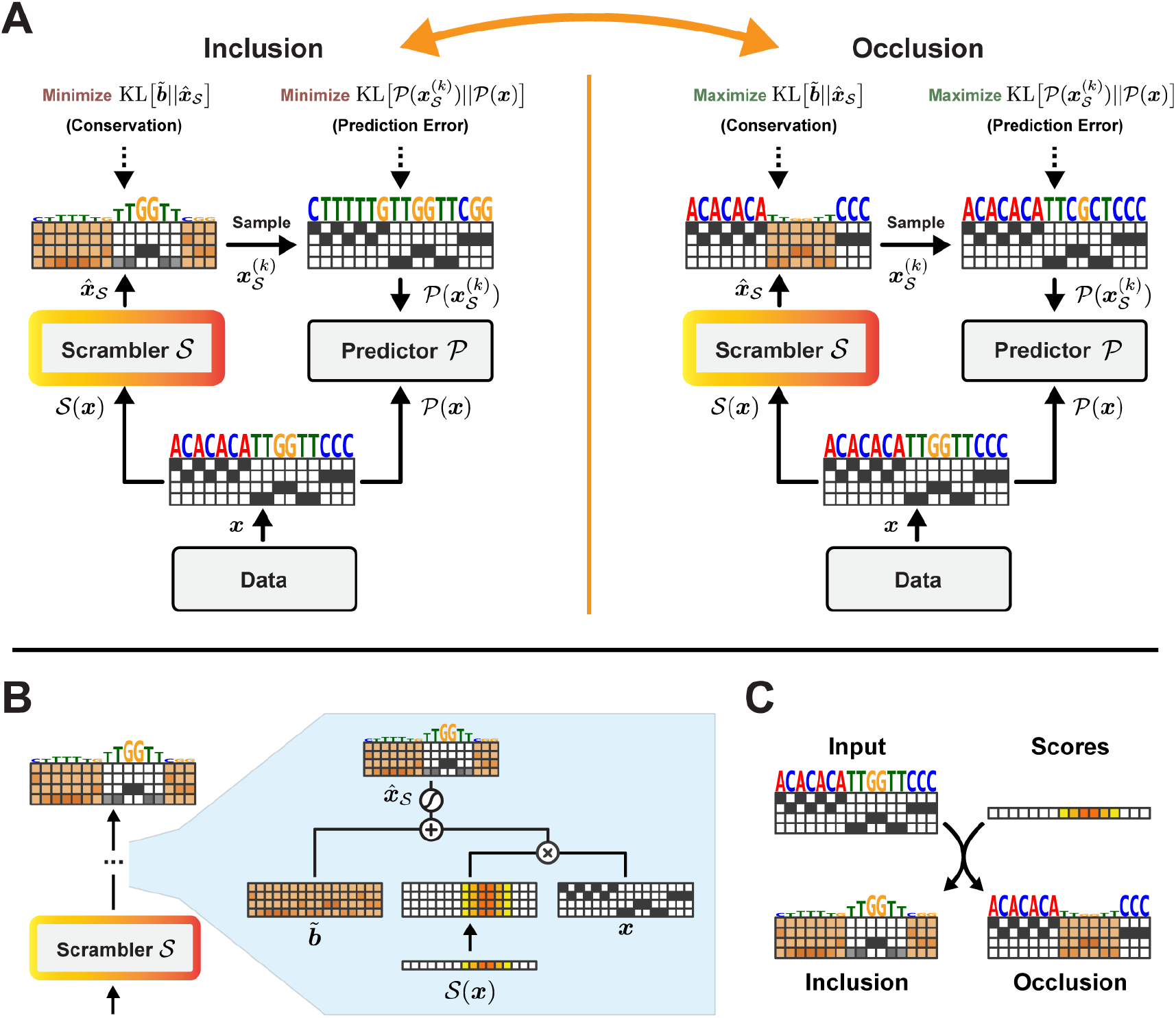
(A) High-level architecture. Inclusion: Maximize the entropy of the PSSM predicted by the Scrambler and minimize the prediction error of samples drawn from it. Occlusion: Minimize PSSM entropy and maximize sample prediction error. (B) The temperature-based masking operation. The Scrambler generates sampling temperatures (‘importance scores’) for the PSSM. (C) The Inclusion and Occlusion scrambling operations. Individual nucleotides (or amino acid residues) are perturbed by raising the sampling temperature of its corresponding categorical feature distribution.

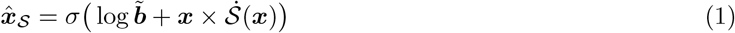

Here, *σ* denotes the softmax 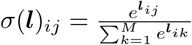 and 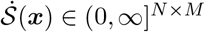 represent the importance scores 𝒮 (***x***) which have been broadcasted at position *i* to all channels *j*. In all experiments, the Scrambler 𝒮 is a residual network of dilated convolutions (He et al., 2016; Figure S1A-C). The output 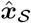 becomes a parameterization of a probability distribution of the input. Specifically, 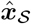 is a set of *N* categorical softmax-nodes, or a PSSM. The role of 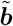 is to keep samples from this PSSM in-distribution and along the manifold of valid patterns, and is here taken as the mean input pattern across the training set (Equation 10). When 𝒮 (***x***)_*i*_ is close to 0, 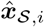 becomes 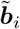 (the background distribution) and when 𝒮 (***x***)_*i*_ is close to ∞, 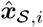 becomes ***x***_*i*_ (the original input). 𝒮 (***x***)_*i*_ thus defines the inverse feature sampling temperature at position *i* in the PSSM.

*K* discrete samples 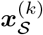 drawn from 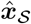 are passed to the predictor 𝒫 and gradients are backpropagated to 𝒮 using either Softmax Straight-Through estimation (Chung et al., 2016) or the Gumbel distribution (Jang et al., 2016). By comparing the predictions 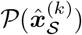 of the scrambled input samples to the original prediction 𝒫 (***x***), we train the Scrambler 𝒮 to minimize a predictive reconstruction error subject to a conservation penalty which enforces high entropy. We refer to this formulation as the *Inclusion*-Scrambler, as it must learn to include features to reconstruct the original prediction while maintaining high entropy of 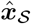. Alternatively, we can train 𝒮 to find the smallest set of features in ***x*** to randomize (i.e., maximizing the conservation of 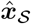) to maximally perturb 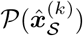 from 𝒫 (***x***) (Figure 1A, right). We refer to this inverse formulation as the *Occlusion-Scrambler*. For both formulations, we define the reconstruction error as the KL-divergence 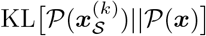 between scrambled and original predictions. To minimize the conservation of 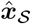 while still keeping samples in-distribution, we optimize the KL-divergence 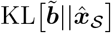 between 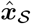 and the background distribution 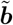. We control the expected entropy by fitting 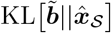 to a target conservation value *t*_bits_ rather than minimizing or maximizing it unbounded. The full training objective for the Inclusion-Scrambler is given in Equation 2.

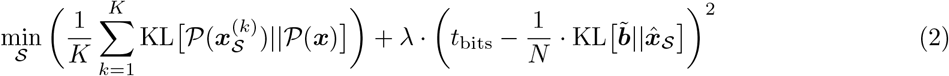

We compare several different attribution methods in our experiments. The baseline method used for comparison, *Perturb*, exchanges the categorical value of one letter or pixel at a time and estimates the absolute value in predicted change as the importance score. Comparisons are made against Perturb (baseline), Gradient Saliency (Simonyan et al., 2013), Guided Backprop (Springenberg et al., 2014), Integrated Gradients (Sundararajan et al., 2017), DeepLIFT (Shrikumar et al., 2017; using RevealCancel for MNIST and Rescale from DeepExplain for the remaining tasks; Ancona et al., 2017), SHAP DeepExplainer (Lundberg et al., 2017) and the preservation/perturbation methods of Fong et al. (2019; ‘TorchRay’), Dabkowski et al. (2017; ‘Saliency Model’) and Carter et al. (2019, ‘SIS’). For comparison, we also use a version of the Scrambler with a zero-based masking operator identical to L2X and INVASE (referred to as ‘Zero Scrambler’). See Methods for a detailed description of how each method was used and which task(s) it was tested on.

### Finding Salient Pixels in MNIST Images

We first used the Scrambler to visualize important regions in the binarized MNIST digits. The predictor was a convolutional neural network (CNN) trained to discriminate between all ten digits (Figure 2A). We trained one Inclusion Scrambler (*t*_bits_ = 0.005, *λ* = 500) and one Occlusion Scrambler (*t*_bits_ = 0.3, *λ* = 500). We performed two benchmark comparisons to other attribution methods (Figure 2B): first, based on the importance scores of each method, we replaced all but the top *X*% most important pixels with random samples from the background and measured the KL-divergence between the predictions of the original and randomized test set images. Next, we inversely tested how well these top-ranked features perturbed the prediction by replacing only the top *X*% pixels with random samples. The Scramblers were superior to other tested methods in each benchmark, as the median KL-divergence was the lowest and highest, respectively. Interestingly, the Scramblers identified the upper open regions of ‘3’ and ‘5’ as important, indicating that the classifier uses the background pixels rather than pixels on the digits when making its prediction.

**Figure 2:**
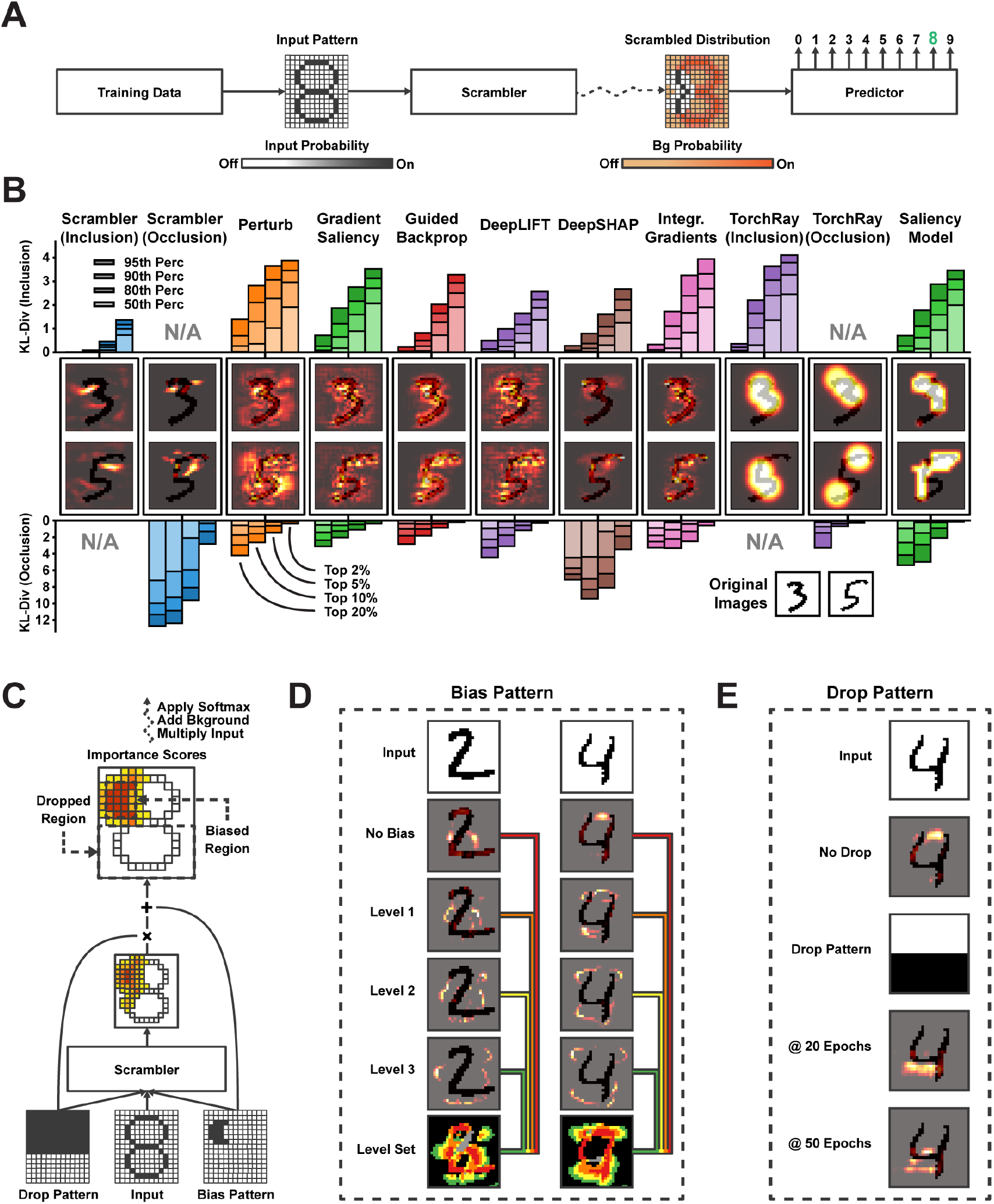
(A) Inclusion and Occlusion-Scrambler setup for MNIST 3 vs. 5 attribution. (B) Attributing feature importance for binarized MNIST digits ‘3’ and ‘5’. (Upper bar chart) Keeping the top *X*% pixels according to importance scores and replacing all other pixels with random values. (Lower bar chart) Replacing the top *X*% pixels with random values. (C) Importance score dropout and bias layers for the MNIST task. (D) Example of dynamically sized feature sets using progressive bias patterns for differentiating digits ‘2’ vs. ‘4’. The pixels found at each level are added to the next level of bias pattern to discover the next set of features. (E) Example of finding alternate salient features when dropping the upper half of the image importance scores.

### Identifying Multiple Salient Feature Sets with Dropout and Bias Layers

There are often multiple feature sets a classifier can use within a single input pattern to make its prediction. Scramblers can be trained to select these alternate salient feature sets dynamically using a dropout pattern ***D*** which is used to mask some positions in the generated scores 𝒮 (***x***). Similarly, bias patterns ***T*** are used to encode specific positions to be included in the solution. Importantly, the Scrambler 𝒮 also receives ***D*** and ***T*** as additional input, allowing the network to learn to output alternate (or additional) scores conditioned on which positions were dropped or enforced. (Figure 2C, Supplementary Fig. S2A-C; see Methods for details). During training, we sample random dropout- and bias patterns. At attribution time, we supply specifically crafted patterns to exclude or enforce in the predicted solution to enable targeted feature attributions.

For the MNIST classifier, bias patterns allow us to explore features beyond the initial salient pixels found by default (Figure 2D). Conversely, we can use dropout patterns to focus attribution to pixels in a subsection of an image to discover new salient features (Figure 2E). In Supplementary Fig. S2D-E, we trained a Scrambler with tunable divergence against the background 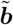, demonstrating how the entropy of the solution can be set dynamically at attribution time, thus providing solutions at multiple scales of *t*_bits_ with a single network.

### The cis-Regulatory Code of Alternative Polyadenylation

Next, we used Scramblers to interpret regulatory sequence elements in the 3’ untranslated region (UTR) of pre-mRNA. Specifically, using APARENT (Bogard et al., 2019) – a CNN capable of predicting cleavage and polyadenylation from primary sequence – we trained an Inclusion-Scrambler to reconstruct isoform predictions for the original APARENT training data (*t*_bits_ = 0.25, *λ* = 1; Figure 3A). As we anticipated important polyadenylation features like RNA binding protein (RBP) recognition motifs to consist of short subsequences, we regularized the Scrambler by fixing the final layer of the network to a Gaussian filter (reminiscent of a technique proposed by Fong et al., 2019) to encourage the selection of contiguous nucleotides for masking. We found that the regularized Scrambler learned to recognize known regulatory binding factors associated with alternative polyadenylation (Supplementary Fig. S3A), such as CFIm, CstF, HNRNPL, ENOX1, PABPN1, HuR, and RBM4 (Di Giammartino et al., 2011; Shi et al., 2012; Elkon et al., 2013; Tian and Manley, 2017). The Scrambler outperformed other attribution methods when reconstructing the isoform predictions using only the top-ranked features (Supplementary Fig. S3B-C). When removing the fixed Gaussian filter from the Scrambler, the network learned patterns that resulted in more extreme perturbations (Supplementary Fig. S3D-E). However, when inspecting these patterns, they clearly exploited the importance of scattered T’s in favor of selecting biologically relevant motifs (Supplementary Fig. S3F).

**Figure 3:**
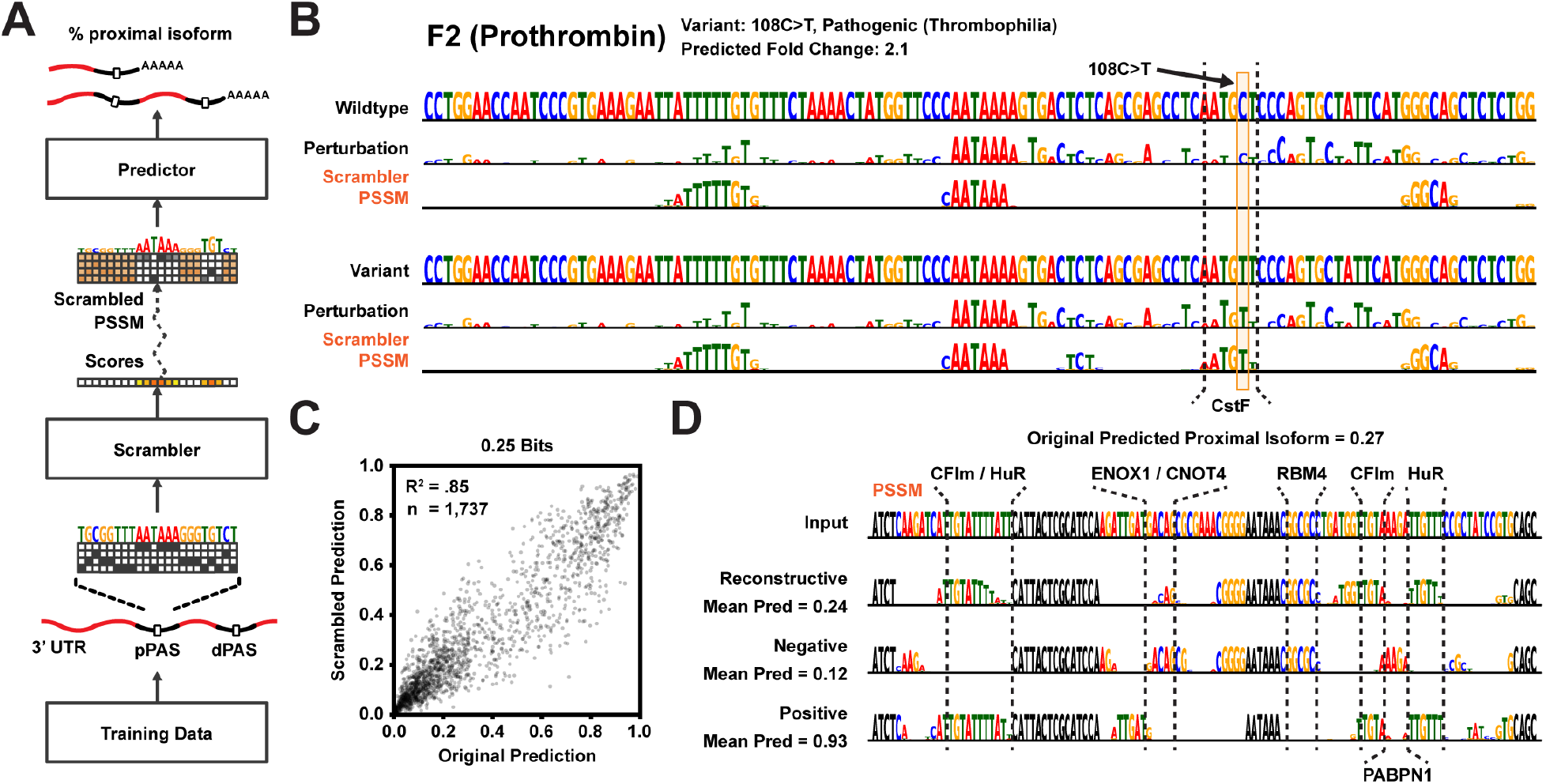
(A) Scrambler architecture for polyadenylation isoform predictions using the pre-trained model APARENT (Bogard et al., 2019) as the predictor. (B) Example attributions of a deleterious variant (*108C>T*) in the 3’ UTR of the Prothrombin (*F2*) gene, comparing the Perturbation method to an Inclusion-Scrambler trained on the APARENT data. The variant creates a cryptic CstF binding site (annotated), increasing polyadenylation efficiency and ultimately steady state mRNA levels. The Scrambler was trained with *t*_bits_ = 0.25 target bits of conservation. (C) APARENT predictions of original test set sequences compared to predictions made on sequence samples produced by the Scrambler. (D) Example of Inclusion-Scramblers trained to reconstruct, maximise or minimize predictions, thus finding overall important, enhancing or repressing motifs, respectively.

### Interpreting Genetic Variants of APA

By running the Scrambler on wild-type and variant human polyadenylation signals, we can interpret the functional impact of mutations. For example, the Scrambler correctly detects the creation of a cryptic CstF binding site caused by a deleterious variant (*108C >T*) in the *F2* (Prothrombin) gene, which is known to cause or contribute to Thrombophilia due to increased polyadenylation efficiency (Figure 3B; Wylenzek et al., 2001; Danckwardt et al., 2004; Takagaki and Manley, 1997). The detected CstF motif is qualitatively better separated from the Scrambler sequence logo’s background compared to Perturbation. Furthermore, the Scrambler appears highly sensitive in detecting loss of RBP binding motifs due to single nucleotide variants for several example human polyadenylation signals (Supplementary Fig. S3G).

### Reconstructive, Enhancing and Repressive Motifs

Interestingly, with as little as 0.25 out of 2.0 target bits of mean conservation in the scrambled sequences, there was very high correlation between scrambled and original sequence predictions from the APARENT test set (*R*^2^ = 0.85; Figure 3C). In other words, a small set of regulatory features in each polyadenylation signal explained most of the variation. Next, we altered the Scrambler training objective to find the smallest set of features that maximize or minimize a prediction rather than reconstructing it (see Methods for details). We found that the Scrambler could partition cis-regulatory elements within polyadenylation signals into enhancing (e.g., CFIm, HuR, Cstf) and repressive motifs (e.g., ENOX1, RBM4, PABPN1), in agreement with the known function for these motifs and associated RBPs (Figure 3D, Supplementary Fig. S3H).

### The Rules of Translation Efficiency in the 5’ UTR

The translation efficiency of mRNA is controlled by complex regulatory logic in its 5’ UTR. For example, an in-frame (IF) start and stop codon create an IF upstream open reading frame (uORF), which typically represses translation, while neither an IF upstream start or upstream stop codon individually represses translation (NAND logic). Sequences with multiple IF starts and stops can further complicate translational logic by creating NAND-OR hybrid functions with overlapping IF uORFs.

We trained Inclusion-Scramblers to reconstruct the mean ribosome Load (MRL – a proxy for translation efficiency) of synthetic 5’ UTRs as predicted by the CNN Optimus 5-Prime (Sample et al., 2019; Figure 4A). To test how well attribution methods detect multiple regulatory elements, we generated synthetic 5’ UTR datasets with one (1 IF start, 1 IF stop), two (1 IF start/2 IF stops or 2 IF starts/1 IF stop), or four (2 IF starts/2 IF stops) overlapping IF uORFs. We then measured to what extent the most important nucleotides of each method corresponded to the IF start and stop positions (Figure 4B, Supplementary Fig. S4B-D). In the simplest case of a single start and stop, all methods had high accuracy for recovering these elements. However, as the number of IF uORFs increased, the Scrambler often outperformed other methods considerably. An example 5’ UTR with multiple IF uORFs is shown in Figure 4C (and Supplementary Fig. S4E); here, the multiple stops effectively hide one another from many local attribution methods.

**Figure 4:**
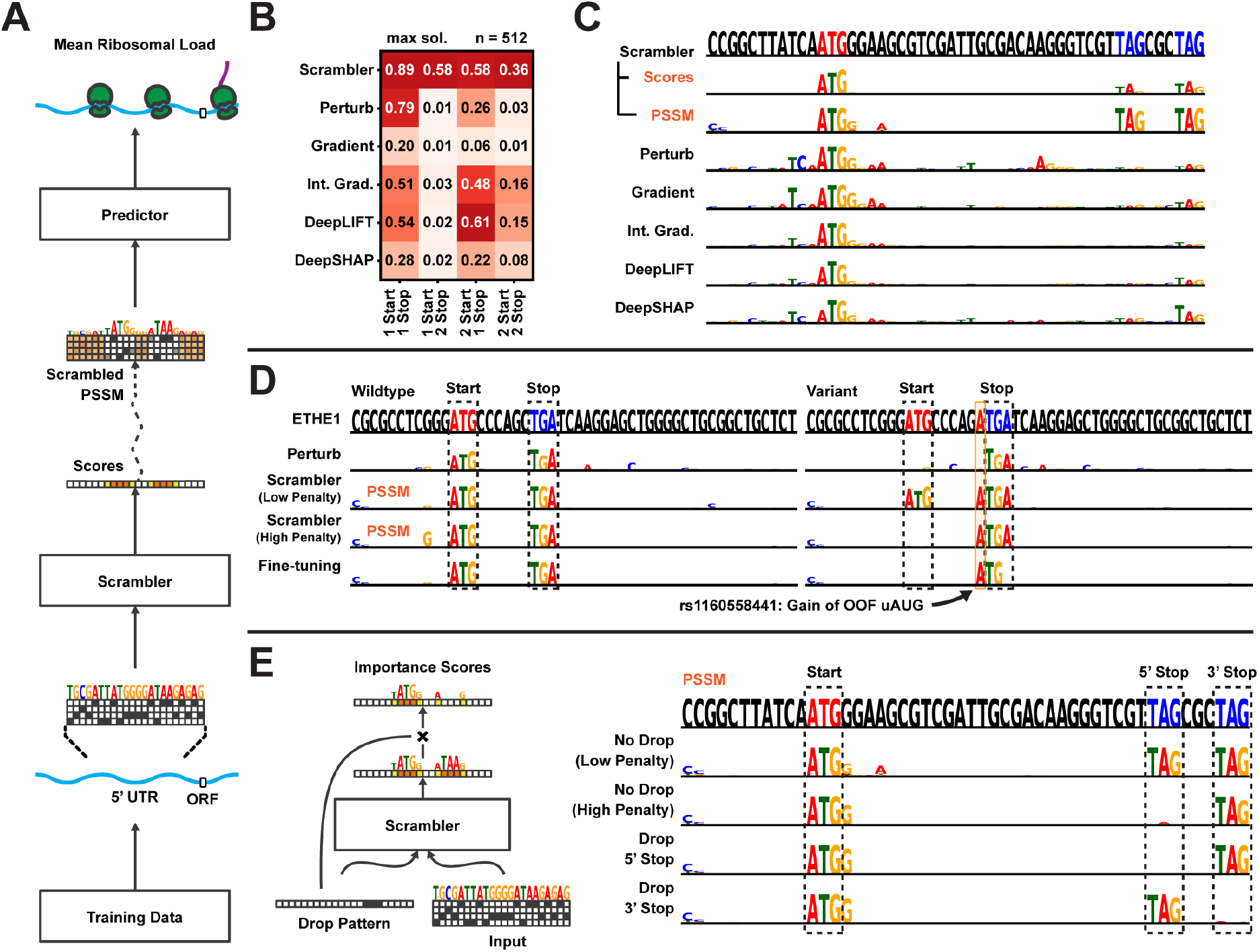
(A) Scrambler architecture for 5’ UTR ribosome load predictions using the pre-trained model Optimus 5’ (Sample et al., 2019) as the predictor. (B) Average recall of in-frame (IF) start and stop positions in a test set of synthetic IF uORF 5’ UTRs. In this test, all start and stop codons must be among the highest scored nucleotides of a sequence to count as successfully recalled for a given method. (C) Example attribution in a 5’ UTR with multiple IF stop codons. The Scrambler was trained with a low entropy penalty (weight = 1, *t*_bits_ = 0.125). (Top sequence) Red letters = IF start codon, Blue letters = IF stop codon. (D) Interpreting a rare, functionally silent mutation (*rs1160558441*) in the 5’ UTR of the *ETHE1* gene, where an out-of-frame (OOF) upstream start codon (uAUG) is created within an in-frame (IF) upstream open reading frame (uORF). Two Scrambler networks were trained, one with a low entropy penalty (weight = 1, *t*_bits_ = 0.125) and one with high penalty (weight = 10, target bits = 0.125). The ‘Fine-tuning’ optimization was performed on the importance scores generated by the Low penalty-Scrambler. (E) Finding alternate IF uORF regions by separately dropping each of the Stop codons.

### Interpreting Genetic Variants in the 5’ UTR

The combinatorial nature of out-of-frame start codons (OOF uAUGs) and IF uORFs makes variant interpretation in human 5’ UTRs complicated (Calvo et al., 2009; Araujo et al., 2012; Whiffin et al., 2020). In Figure 4D, we interpret a variant, *rs1160558441*, which creates an OOF uAUG that Optimus 5-Prime predicts to be functionally silent. The OOF uAUG is created within an IF uORF, which hides its effect, as either element alone is sufficient to repress translation. Therefore, the variant sequence is not interpretable by Perturbation as the uAUGs are not found. A Scrambler trained with a low entropy penalty (*t*_bits_ = 0.125, *λ* = 1) detects both possible interpretations. With a higher penalty (*λ* = 10), the Scrambler must find a smaller set of salient features and marks only the OOF uAUG as important, which is sufficient to explain the repression. However, rather than just identifying ‘ATG,’ the Scrambler identifies the subsequence ‘ATGA,’ even though the trailing adenine is not important for the explanation. This occurs because the number of salient features is highly variable in any given 5’ UTR, and so a Scrambler trained for a target entropy *t*_bits_ may ‘over-interpret’ some examples.

### Per-example Fine-tuning Removes Interpreter Artifacts

To overcome issues with over-interpretation for datasets with a highly variable number of important features, we apply a per-example fine-tuning step: starting with 𝒮 (***x***), we optimize new scores ***s*** by gradient descent which are specific to ***x*** and remove excessive features of 𝒮 (***x***) and maximize the entropy of the PSSM unbounded. Specifically, ***s*** is optimized to subtract scores from 𝒮 (***x***) in order to maximize the PSSM entropy 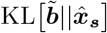 unbounded, while still minimizing the predictive reconstruction error 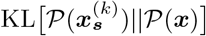. As can be seen for the variant example (Figure 4D), fine-tuning of the Low entropy penalty Scrambler removes the trailing ‘A’. We also compared the Scrambler with and without fine-tuning to interpret silent uORF-altering and gain-of-function mutations (Supplementary Fig. S4G). Importantly, when compared on one of the benchmarks of Figure 4B, we find that applying per-example fine-tuning to the pre-trained Scrambler scores produces more robust attribution results than individual per-example optimization of importance scores (Supplementary Fig. S4H). This is likely because the latter approach can become stuck in local minima and produce poorly generalizable interpretations, essentially ‘overfitting’ to spurious predictor signals.

### Dropout for Alternate Open Reading Frame Discovery

Sequences with multiple IF uORFs are examples of patterns with redundant salient feature sets. By adding an importance score dropout layer to the Inclusion-Scrambler and training it on random dropout patterns, we obtain a model capable of dynamically proposing different IF uORFs as attributions (Figure 4E, Supplementary Fig. S4F, I). Without dropout and a low entropy penalty (*t*_bits_ = 0.125, *λ* = 1), the Scrambler marks both IF stops (i.e. both IF uORFs) as important in the example from Figure 4C. However, with a higher penalty (*λ* = 10), the Scrambler finds a smaller interpretation with only one IF stop. When using dropout patterns to exclude either of the IF stops, the Scrambler dynamically finds the alternate IF uORF.

### The Determinants of Protein-Protein Interactions

Here we apply Scramblers to interpret predicted interactions for pairs of proteins. This can be challenging, as proteins are defined along a narrow manifold of stably folded sequences. Any interpretation method must ensure that masked or perturbed sequences stay in distribution for predictions to remain accurate. We focused on a set of rationally designed coiled-coil heterodimers, where binding specificity is induced by hydrogen bond networks (HBNets) at the dimer interface (Maguire et al., 2018; Chen et al., 2019). Using approximately 180, 000 designed heterodimers as positive training data and randomly paired monomers as a negative set, we trained a recurrent neural network (RNN) to predict dimer binding (Figure 5A, Supplementary Fig. S5A; AUC = 0.96 on held-out test data). These coiled-coil proteins follow a conserved heptad repeat structure, but binder sequences of different lengths have their heptad motifs shifted by different offsets. Consequently, when training Scrambler networks on this data, we used a separate background distribution for each sequence length. A test set of 478 designed heterodimers was used to investigate how different attribution methods could capture HBNet positions and residues important for complex stability.

**Figure 5:**
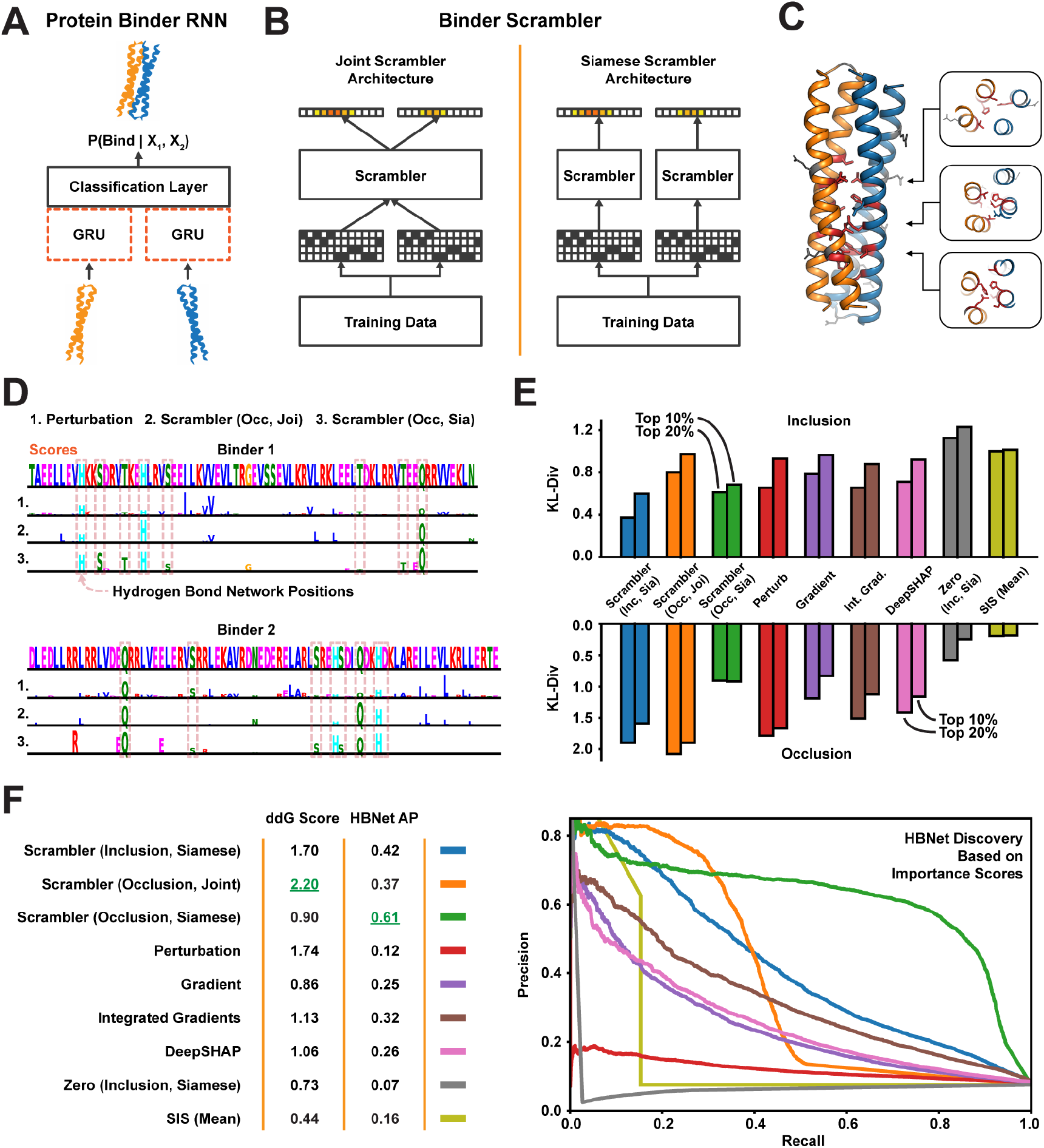
(A) Protein binding predictor. A RNN based on a siamese GRU layer (orange dashed lines). (B) The *joint* and *siamese* Scrambler architectures for protein binder attribution. (C) 3D-visualization of an example binder pair structure. Discovered HBNet residues (by a siamese Occlusion-Scrambler) are shown in red licorice, important residues not part of the HB-net are shown as gray sticks. A cross-section of each of the three buried hydrogen bond networks is shown on the right. (D) Example attribution. Hydrogen bonds at the designed binding interface are marked with dashed red boxes. (E) Keeping the top *X*% residues according to the importance scores of each method and replacing the rest with random amino acids (top), or replacing the top *X*% with random amino acids (bottom), measuring prediction KL-divergence. ‘Zero’ refers to a Scrambler with a zero-based masking operator. (F) Left: *In silico* Alanine scanning benchmark. The 8 most important residues of each method were replaced with Alanines, measuring ΔΔ*G* in Rosetta Energy Units (REU). Shown are the mean ΔΔ*G* differences in a permutation test of 10,000 re-labelings. Right: Precision-recall curves and Average Precision (AP) for discovering the HBNet positions, based on the importance scores of each benchmarked method.

Unlike for the DNA attribution tasks above, each dimerization prediction involves two input sequences. Consequently, there are multiple ways in which to define the Scrambler architecture. We first tested a *joint* Occlusion-Scrambler (Figure 5B; left), which receives both sequences as input and can learn to recognize partner-specific binding features. After training, this architecture identified a subset of HBNet positions and hydrophobic residues at the interface necessary for binding specifically to the cognate partner (Figure 5D). We also tested a *siamese* architecture, which saw only one input sequence at a time and was therefore constrained to learn global, binding partner-independent features (5B; right). This architecture learned to identify nearly all HBNets, as these are the best determinants of *binding specificity* to any possible partner (Figure 5C-D). We compared these Scrambler architectures to other attribution methods based on how much the positions with largest absolute-valued score either preserved or perturbed the RNN predictions (Figure 5E; *t*_bits_ = 0.25, Inclusion; *t*_bits_ = 2.4, Occlusion). The siamese Inclusion-Scrambler and joint Occlusion-Scrambler had the best (lowest and highest) median KL-divergence for these tasks.

### Importance Scores Reflect Dimer Complex Stability and Hydrogen Bonds

Next, using in-silico Alanine scanning, we estimated the mean difference in ΔΔ*G* between the original and mutated dimers for the eight most important residues per binder and attribution method (Ford et al., 2020). We also compared the methods by their ability to discover HBNet positions based on their importance scores. We found that the joint Occlusion-Scrambler identified residues that destabilized the complex the most (mean difference ΔΔ*G* = 2.20, Figure 5F), while the siamese model discovered significantly more HBNet positions (AUPRC = 0.61). These results support the idea that the siamese architecture learned discriminative features for both interacting and non-interacting pairs – HBNet residues – while the joint model learned features important for positive interaction, such as hydrophobic residues at the interface. While Integrated Gradients and DeepSHAP perturbed the RNN predictions more when considering only their positive-valued scores (Supplementary Fig. S5B, left), the Scramblers had better ΔΔ*G* scores and HBNet discovery rates (Supplementary Fig. S5B, right). This suggests that using a generative model results in more generalizable features by overcoming spurious RNN signals. We performed additional comparisons to other attribution methods in Supplementary Fig. S5C-D and **??**. We found that the temperature-based masking operator of Equation 1 consistently outperforms other masks and that generative interpretations were better than per-example masking approaches.

Finally, we tested the Scrambler against different versions of the ‘hot-deck’ SIS method (Supplementary Fig. S5E; Carter et al., 2019; Carter et al., 2020): For each iteration of SIS, we sampled a number of background sequences and used these to mask de-selected features. The mean sample prediction was used as the function value. Similarly, we varied the number of Gumbel patterns sampled from the Scrambler PSSM during training (*K* in Equation 2). The Scrambler operated well with few (≥ 4) samples and consistently outperformed SIS with 32 samples, both in predictive reconstruction and ground-truth comparisons. Additionally, the total number of predictor queries required to interpret the entire dataset with comparable quality was ∼ 70 times lower than SIS. Interestingly, using a simple masking scheme such as mean features resulted in bad interpretations for the dimer RNN predictor.

### Interpreting Protein Structure Predictions

Recent advances in deep learning have made homology-free protein tertiary structure prediction possible. Such neural networks use the primary sequence and corresponding multiple sequence alignment (MSA) as input to predict three-dimensional atom-atom distances and backbone torsion angles (Senior et al., 2020). Here, we apply Scramblers to the predictor trRosetta (Yang et al., 2020) to detect important sequence features for predicted structures. Since it is computationally heavy to query the predictor – and would likely require many iterations to train a Scrambler network – we demonstrate the approach for a single protein at a time (Figure 6A). A trainable vector of scrambler scores (PSSM temperatures) is used to perturb the input sequence and MSA to minimize or maximize the total KL-divergence between predicted contact distributions (see Methods for details).We tested our approach on one of the alpha-helical hairpin binders from the previous dimer data set (Figure 6B). PSSMs for the hairpin binder were optimized for the Inclusion (Figure 6C) and Occlusion objectives (Figure 6D). Both PSSMs used hydrophobic leucines and a symmetry-breaking glycine in the hairpin region to reconstruct or distort the original hairpin structure prediction, which aligns well with previous results (Chen et al., 2019). We further tested the method on naturally occurring proteins with more complex 3D structures (Supplementary Fig. S6A-B).

**Figure 6:**
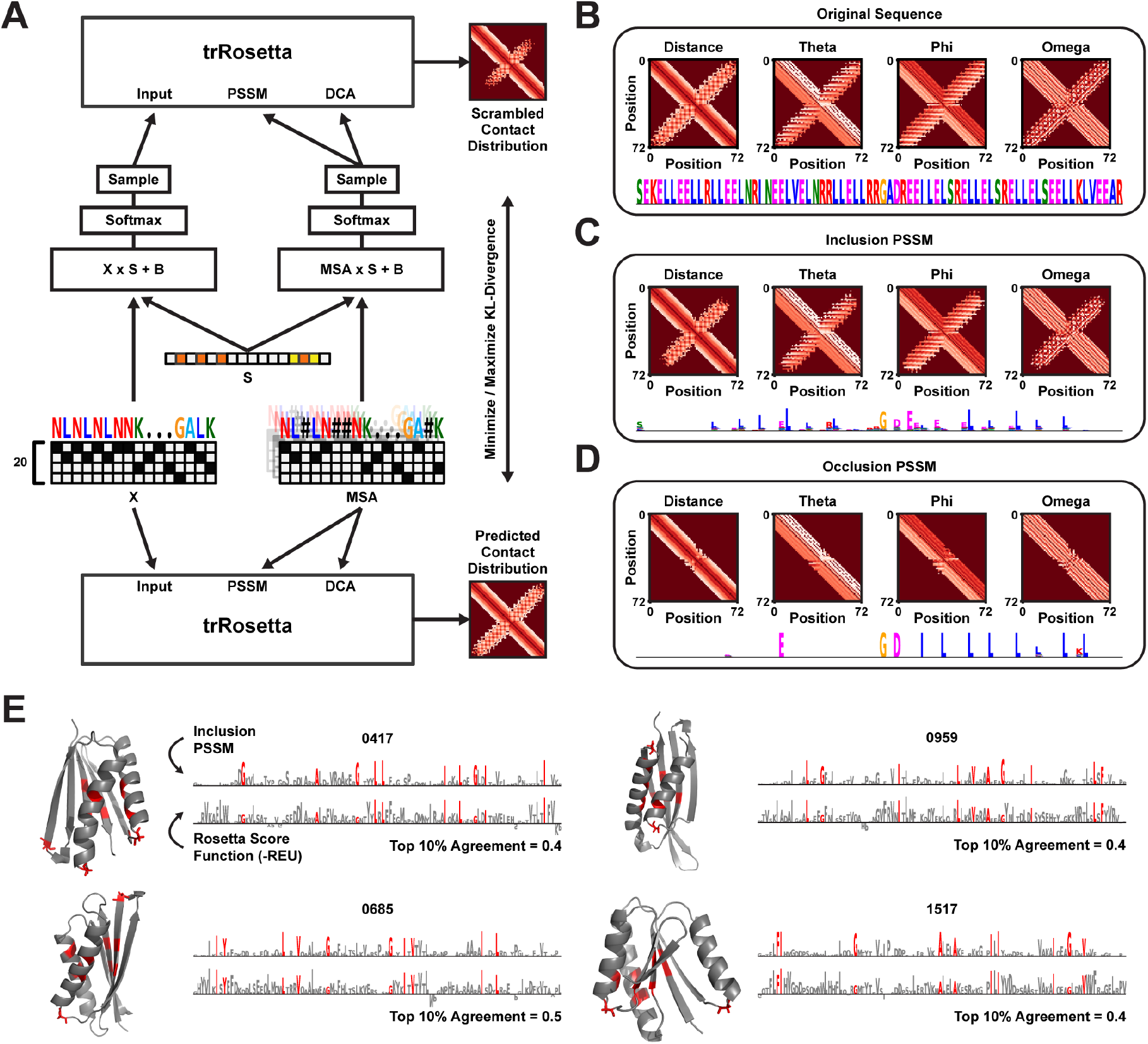
(A) The single-sequence Scrambling architecture for trRosetta. Both the primary sequence (‘X’) and the multiple sequence alignment (‘MSA’) are scrambled with the same trainable mask (‘S’). (B) Alpha-helical hairpin protein sequence and original structure prediction. (C) Inclusion-Scrambled PSSM of the hairpin protein, with the average predicted distance- and angle distributions for 512 samples drawn from the PSSM. (D) Occlusion-Scrambled PSSM of the hairpin protein and average structure prediction for 512 samples. (E) Inclusion-Scrambled PSSMs of four *de novo* proteins which have been designed by deep network hallucination (Anishchenko et al., 2020). Each PSSM was optimized for *t*_bits_ = 1.0 target bits of conservation, using the amino acid frequency of naturally occurring sequences from the Protein Data Bank as background distribution. The bottom sequence logos represent the Rosetta score function breakdown per residue (*−*REU). The agreement between the 10 most important residues according to the importance scores and the absolute energy values ranged between 0.4 and 0.5. The 10 most important residues according to the Scrambler scores are colored red (both in the sequence logos and 3D illustrations).

Next, we altered the Scrambling architecture to enable MSA-free interpretation of *de novo* engineered proteins lacking natural sequence homology (Supplementary Fig. S6C). We optimized Inclusion-PSSMs to reconstruct the structure prediction of four proteins designed by deep network hallucination (Figure 6E; Anishchenko et al., 2020). Interestingly, important features found by the Scrambler included hydrophobic residues in the core for all four proteins (10 most important residues colored red). For validation, each protein’s structure prediction was relaxed, and the per-residue breakdown of the Rosetta score function was computed (Alford et al., 2017). We found that the per-residue energy values agreed quite well with the Scrambler importance scores (mean top 10 residue agreement = 0.425). However, note that some features marked by the Scrambler as important for the structure prediction are missed by the Rosetta energy breakdown. For example, as can be seen in Figure 6E, the Scrambler marked loop glycines between alpha-helices as important for the structure prediction, even though they do not have a large energetic contribution to overall stability. Glycines frequently occur in loops and turns and are thought to be important for maintaining protein loop structure due to their size and flexibility compared to other residues (Parrini et al, 2005, Krieger, Moglich, and Kiefhaber 2005). The MSA-free approach was also tested on a hairpin-like protein engineered by Activation Maximization (Supplementary Fig. S6D-E; Linder et al., 2020).

## Discussion

Here we introduced a new attribution method – Scrambler Networks – that builds on and extends earlier work in input masking approaches. The goal is to explain an input pattern for a machine learning model by finding the smallest set of features which either reconstruct or destroy its prediction (Fong et al., 2017; Dabkowski et al., 2017; Chen et al., 2018; Yoon et al., 2018; Fong et al., 2019; Carter et al., 2019). Scrambler networks are based on learning a generative model that predicts a set of real-valued importance scores given an input pattern. These predicted scores are used as sampling temperatures within a PSSM that interpolates between the current input and a non-informative background distribution. We demonstrated the performance of Scramblers on several attribution tasks, ranging from identifying cis-regulatory elements in RNA sequences to protein-protein interactions and protein tertiary structure. Furthermore, Scramblers can be trained to find not only reconstructive, but also enhancing and repressive features, and can be used to dynamically find multiple salient feature sets within a single input pattern. For prediction problems where the salient feature sets vary widely in size, per-example fine-tuning can be employed to correct positions that were misidentified as important.

At the core of mask-based interpretation is the masking operator, which must be carefully chosen according to the problem domain. Earlier work in computer vision masked input patterns by fading or blurring (Fong et al., 2017; Dabkowski et al., 2017; Fong et al., 2019), and other methods for discrete variable selection masked features with zeros (Chen et al., 2018; Yoon et al., 2018). Neither of these operators are suitable for predictors trained on biological sequences, which expect discrete one-hot encoded patterns as input and predict poorly on patterns outside of the training data manifold. The Scrambler masking operator is conceptually similar to ‘hot-deck’ masking proposed in the *SIS* method by Carter et al. (2019) and in the interpretation method of Zintgraf et al. (2017), where deactivated features are replaced with random samples from the marginal distribution of that input. Scramblers generalize this concept by maintaining a categorical feature sampling distribution at each position in the sequence, where the predicted temperature (inverse importance score) defines how random a sample from that position will be on a continuous scale. An immediate benefit is that, in contrast to SIS, Scramblers generate real-valued importance scores that reflect an internal ranking of feature importance. This masking operator also shares similarities with the counterfactual masking introduced by Chang et al. (2018), but the temperature-based operator is simpler as it does not require training a generative fill-in model.

We showed in extensive benchmarks that temperature-based masking outperforms the hot-deck masking done with SIS, as well as any of the zero-, mean- or sample-masking schemes which are used in *L2X* (Chen et al., 2018) and *INVASE* (Yoon et al., 2018). By training a parametric generative model, we end up with a more sample-efficient method at interpreting patterns than per-example methods (Fong et al., 2017; Carter et al., 2019), as measured by the cumulative number of queries made to the predictor. Also, by optimizing over a large training set, Scramblers avoid overfitting to spurious per-example signals and learn to generalize feature importance. Generalizable attributions are highly desirable in genomics and proteomics as they allow us to infer higher-quality biological discoveries from trained predictors, thereby offering richer insights into molecular processes that were previously considered black boxes. We envision that Scramblers will be particularly useful for rational sequence design, where they can be employed to either validate known design rules (as demonstrated on the protein-protein interaction task) or – as demonstrated on the protein structure task – to discover determinants which data-driven design methods have produced.

## Methods

### Scrambling Neural Network Definition Masking Operator

Let ***x*** ∈ {0, 1}^*N ×M*^ be a one-hot-encoded sequence and let 𝒮 (***x***) ∈ (0, ∞]^*N*^ be the corresponding real-valued importance scores predicted by Scrambler network 𝒮. The channel-broadcasted importance scores 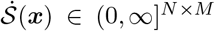 are defined as 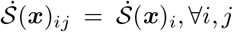. 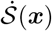 are used as (differentiable) interpolation coefficients in log-space between background distribution 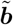 and the current input pattern ***x*** according to Equation 1 for the Inclusion objective. Conversely, the scrambling operation for the Occlusion-model is defined in Equation 3 below. Here, the expression 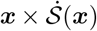 has been replaced with 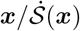. Either of these formulas return a softmax-relaxed distribution (a Position-Specific Scoring Matrix; PSSM) 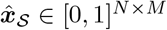, which can be interpreted as a representation of the input sequence where the information content at position *i* has been perturbed according to the sampling temperature 𝒮 (***x***)_*i*_ generated by the Scrambler.

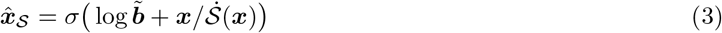

Given the scrambled PSSM 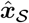, we sample approximate discrete one-hot-coded patterns 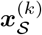 using the Concrete distribution (or Gumbel-distribution; Jang et al., 2016), enabling differentiation:

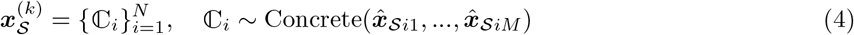

Alternatively, we sample exact discrete patterns 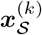 from 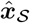 and use the Softmax Straight-Through Estimator (ST-sampling; Chung et al., 2016) to approximate the gradient 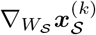 of the input pattern with respect to the Scrambler network weights. To reduce variance during training, we draw *K* samples 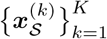 at each iteration of gradient descent and walk down the average gradient (by default, *K* = 32).

### Objective Functions

Given the scrambled PSSM 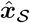, a collection of differentiable sequence samples 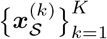 and the background distribution 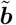, the Scrambler is trained by backpropagation to either minimize Equation 2 (Inclusion) or Equation 5 below (Occlusion). For predictor models 𝒫 with a sigmoid or softmax output (classification), the prediction reconstruction error between 𝒫 (***x***) and 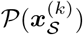 is the standard cross-entropy error. For regression tasks, the cost is defined as the mean squared error. The cost parameter *t*_bits_ defines the target per-position PSSM conservation and *λ* defines the conservation penalty weight. With a smaller value of *λ*, the Scrambler is allowed to generate PSSMs with a larger spread of conservation around the target *t*_bits_ (example: for one input pattern, the scrambled PSSM may have a mean conservation of 0.05, for another pattern, it may be 0.25). With a larger *λ*, the conservation values are forced closer to *t*_bits_.

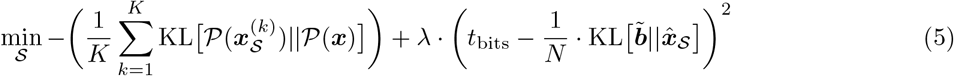

Rather than optimizing the Scrambler to reconstruct predictions using a small feature set, we can train the model to find feature sets that either positively or negatively influence the prediction. For classification tasks, we either maximize or minimize the log-probaility 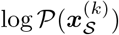 (Equation 6-7). For regression tasks, we either maximize or minimize the regressor 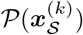 (Equation 8-9).

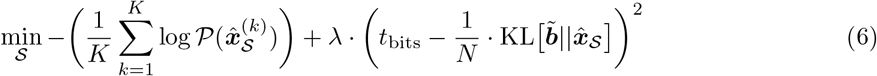

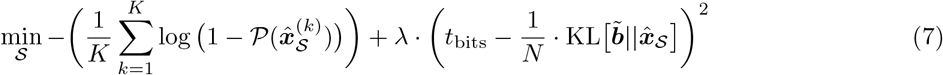

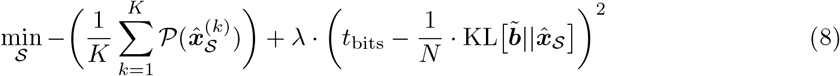

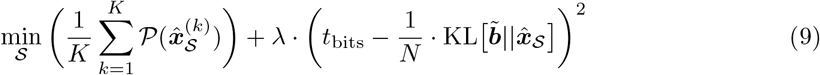

Note that we could have defined the conservation penalty of Equation 2-5 and 6-9 in many other ways. The KL-divergence 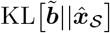 corresponds to the generalized entropy of 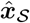 and is particularly suitable for the Inclusion-Scrambler: the expression reaches 0 when 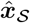 is maximally entropic (with respect to 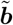). For the Occlusion-Scrambler where entropy should be minimized, it may seem more appropriate to minimize the KL-divergence 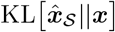 to the original input pattern rather than maximizing 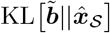, as this cost corresponds to the cross-entropy of 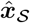 with respect to ***x*** and is 0 at minimum entropy. Alternatively, for both Scramblers, we could simply minimize the area scrambled by the importance scores, *σ*(𝒮 (***x***)) × 2 − 1 (we refer to these as *lum* values). Overall, 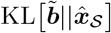 resulted in better performance, which we hypothesize has to do with the information content of 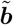 otherwise being lost.

### Background Distribution

The background distribution 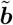 is intended to keep the scrambled sequence samples in-distribution and along the manifold of valid patterns, no matter where on the interpolation line between 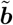 and ***x*** the scrambled distribution 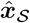 is. There are several possible choices of 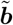, with varying degrees of complexity:

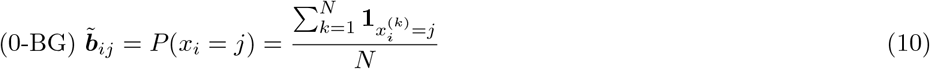

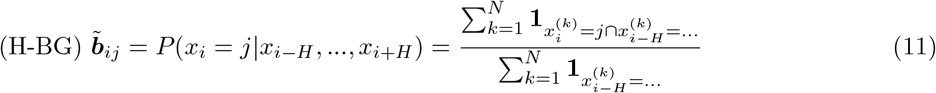

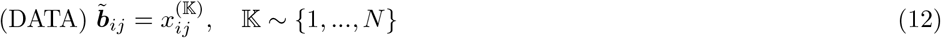

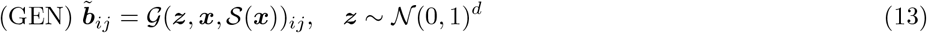

The simplest background distribution, 0-BG, corresponds to the mean input pattern across the training set (i.e. marginal feature probabilities). A more complex background, H-BG, is the distribution of a particular feature conditioned on its neighboring features (Zintgraf et al., 2017). Finally, the DATA- and GEN distributions consist of input patterns randomly chosen from the dataset (DATA), or patterns generated by a distribution-capturing model such as a conditional GAN or Variational Autoencoder (GEN; Chang et al., 2018). In all of our experiments, we relied on the simplest background distribution, 0-BG, since all tested sequence-predictive models behaved well on such samples. Still, for more narrowly defined manifolds, H-BG or DATA/GEN may be more appropriate to stay in-distribution.

### Network Architecture

The Scrambler is based on a Residual Neural Network architecture with dilated convolutions (He et al., 2016; Supplementary Fig. S1A-C). For the MNIST, 5’ UTR and protein binder attribution tasks, the network consisted of 5 groups of 4 dilated residual blocks (20 blocks in total), with filter width 3. For the alternative polyadenylation task, the network had just one group of 4 residual blocks, with filter width 8.

### Mask Dropout and Biasing

Scramblers can find multiple salient feature set solutions for an input pattern using input-masking layers such as dropout- or biasing layers, which either disallow or enforce specific sequence positions in the retained feature set. Specifically, we let ***D, T*** ∈ {0, 1}^*N*^ be the dropout- and biasing masks and apply them to the importance scores by element-wise multiplication and addition respectively:

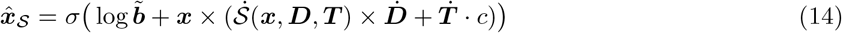

Here *c* is the temperature of the biasing mask ***T*** (smaller *c* allows more randomness in selecting the letter at a biased position, a larger *c* makes it more likely to select the current input letter at the biased position). The Scrambler network S also receives ***D*** and ***T*** as additional input, allowing the network to learn to output alternate scores conditioned on which positions were dropped or enforced. During training, we use random samples of ***D*** and ***T*** (various training schemes are illustrated in Supplementary Fig. S2A-C; for the MNIST task, we used all three schemes; for the 5’ UTR task we used only the first scheme, i.e. uniform random training patterns consisting of squares). After training on randomized dropout and biasing patterns, we can provide hand-crafted patterns at inference time to detect alternate feature sets of new input examples.

### Fine-tuning and Per-example Optimization

For heterogeneous datasets where the number of important features per example is highly variable, we apply a per-example fine-tuning step to the importance scores generated by the (pre-trained) Scrambler network 𝒮 to remove any excessive (or redundant) features. Specifically, for each input pattern ***x***, we generate Scrambler scores 𝒮 (***x***) and optimize the *subtraction scores* ***s*** according to Equation 15-16 (for Inclusion) or Equation 17-18 (for Occlusion) below. The scrambling equation is reformulated with the expression 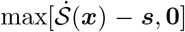; this forces the resulting fine-tuned importance scores to select a subset of the features identified by the original scores 𝒮 (***x***), i.e. the new scores cannot find a new alternate explanation for example ***x***. This restriction prevents ‘overfitting’ to the predictor, which is otherwise a problem with per-example optimization. Architecturally, we do not optimize ***s*** directly; we optimize parameters ***w***, which are instance-normalized and softplus-transformed (***s*** = Softplus(IN(***w***))). In our experiments, we optimize ***w*** for 300 iterations of gradient descent (Adam, learning rate = 0.01).

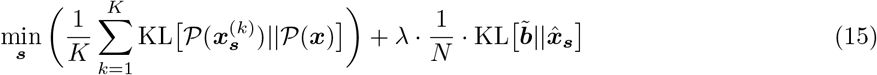

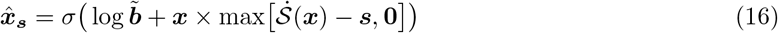

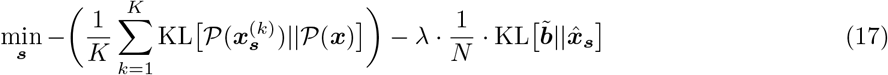

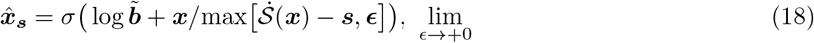

For comparison, we also perform per-example optimization from scratch (i.e., we start with randomly initialized values rather than pre-trained Scrambler scores). To this end, we optimize Equation 15 (for Inclusion) or Equation 17 (for Occlusion). The perturbation operators are identical to Equation 1 and 3, but with individual scores ***s*** per pattern 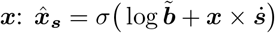 (Inclusion), 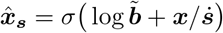 (Occlusion).

### Attribution Tasks and Predictors

#### MNIST

The predictor was a CNN with 2 convolutional layers, 1 max-pool layer and a single fully connected hidden layer. The network was trained for this study. Predicts a 10-way classification score 𝒫 (***x***) ∈ ℝ ^10^ (softmax) given a binarized MNIST image ***x*** ∈ {0, 1}^28*×*28^ as input.

#### APA (3’ UTR)

(Bogard et al., 2019) The predictor was a pre-trained CNN (APARENT) which predicts relative alternative polyadenylation isoform abundance 𝒫 (***x***) ∈ [0, 1] given a 206 nt one-hot DNA sequence ***x*** ∈ {0, 1}^206*×*4^ as input. The trained model was downloaded from: https://github.com/johli/aparent/tree/master/saved_models.

#### Translation (5’ UTR)

(Sample et al., 2019) The predictor was a CNN (Optimus 5-Prime) which predicts mean ribosome load P(***x***) ∈ ℝ given a 50 nt one-hot DNA sequence ***x*** ∈ {0, 1}^50*×*4^ as input. The model was re-trained for this study but the training procedure was based on the code from: https://github.com/pjsample/human_5utr_modeling.

#### Protein heterodimer binders

The predictor was a RNN with siamese recurrent GRU layers, a dropout layer and a single fully connected hidden layer. The network was trained for this study. Predicts protein dimer binding probability 𝒫 (***x***) ∈ [0, 1] (sigmoid activation) given two right-padded, 81 residues long protein binder sequences ***x***_1_, ***x***_2_ ∈ {0, 1}^81*×*20^ as input. Note: When training Scrambler networks on these data, we used a separate background distribution 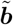 for each unique protein sequence length.

#### Protein structure

(Yang et al., 2020) Predicts distance and angle distributions of backbone atoms in 3D given an input primary sequence. The trained model was downloaded from: https://files.ipd.uw.edu/pub/trRosetta/model2019_07.tar.bz2.

Given a one-hot-encoded sequence pattern ***x*** ∈ {0, 1}^*N ×*20^ and Multiple Sequence Alignment **MSA** ∈ {0, 1}^*K×N ×*21^, trRosetta predicts the atom-atom distance distribution ***D***(***x*, MSA**) ∈ [0, 1]^*N×N×*37^ and backbone torsion angle distributions ***θ***(***x*, MSA**), ***ω***(***x*, MSA**) ∈ [0, 1]^*N×N×*24^, ***ϕ***(***x*, MSA**) ∈ [0, 1]^*N×N×*12^. In the paper, we performed per-example scrambling to interpret example protein structures, i.e. we did not train a generative Scrambler model to predict importance scores. Instead, this method optimizes a set of importance scores ***s*** for each pattern ***x*** that we wish to interpret. In the context of inclusion-scrambling, we find the smallest set of residues that minimizes the total KL-divergence of the target structure:

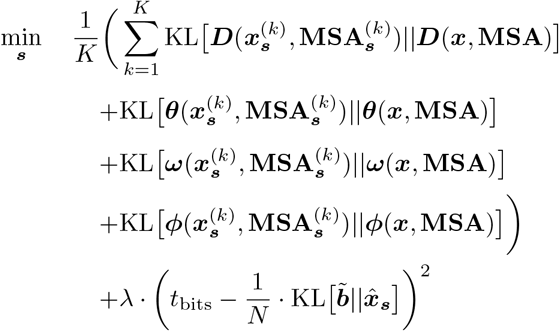

For occlusion-scrambling, we optimize the inverse objective:

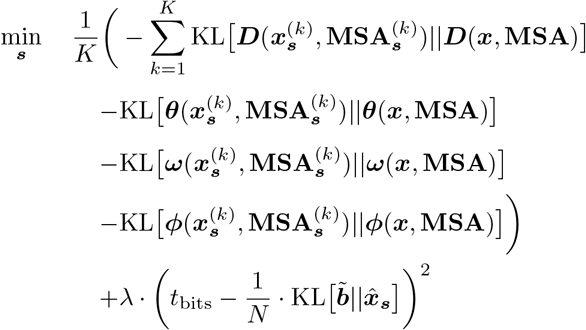

In the above formulations, we define 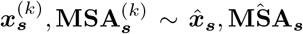 (letter-by-letter Gumbel-sampling). For inclusion-scrambling, we define:

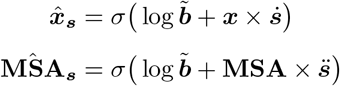

For occlusion-scrambling, we define:

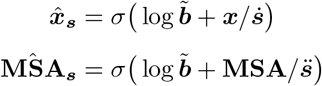

Here 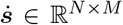 is a channel-broadcasted copy of the importance scores ***s*** ∈ ℝ^*N*^ and 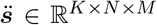 is a MSA-broadcasted copy. We obtain ***s*** from a vector of real-valued (trainable) numbers that we instance-normalize and apply the softplus activation on. We train ***s*** until convergence using the Adam optimizer with learning rate = 0.01, *β*_1_ = 0.5 and *β*_2_ = 0.9.

### Attribution Methods

#### Perturbation

Baseline attribution method where each nucleotide or residue in the input pattern ***x*** is systematically exchanged for every other possible letter, measuring the change in prediction as a proxy for the importance score. Let 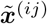 be the sequence pattern corresponding to exchanging the current nucleotide/residue at position *i* for letter *j*. We then calculate the importance score *s*_*i*_ at position i as the absolute mean change in prediction across all letters 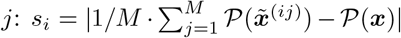. For classification tasks, we compare the log-scores of the predicted probabilities: 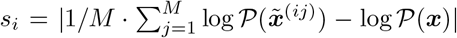.Tested for prediction tasks: MNIST (Figure 2), APA (Figure 3), 5’ UTR (Figure 4) and PPI (Figure 5).

#### Gradient Saliency

(Simonyan et al., 2013) The Gradient Saliency method was executed using the implementation from Ancona et al., (2017). We used either the absolute value of the saliency scores (‘Gradient Saliency’) or the signed scores (‘Gradient Saliency (Signed)’). Tested for prediction tasks: MNIST (Figure 2), APA (Figure 3), 5’ UTR (Figure 4) and PPI (Figure 5).

Code: https://github.com/marcoancona/DeepExplain.

#### Guided Backprop

(Springenberg et al., 2014) The Guided Backprop method was executed using the implementation from Shrikumar et al., (2017). We used either the absolute value of the saliency scores (‘Guided Backprop’) or the signed scores (‘Guided Backpropy (Signed)’). Tested for prediction tasks: MNIST (Figure 2) (the other predictors had compatibility issues).

Code: https://github.com/kundajelab/deeplift.

#### Integrated Gradients

(Sundararajan et al., 2017) Integrated Gradients was executed using the implementation from Ancona et al., (2018). 10 integration steps was used per pattern. The reference was set to the mean sequence pattern (PSSM) of the data set. We used either the absolute value of the saliency scores (‘Integrated Gradient’) or the signed scores (‘Integrated Gradients (Signed)’). Tested for prediction tasks: MNIST (Figure 2), APA (Figure 3), 5’ UTR (Figure 4) and PPI (Figure 5).

Code: https://github.com/marcoancona/DeepExplain.

#### DeepLIFT

(Shrikumar et al., 2017) For the MNIST task, the RevealCancel method was used. For all other tasks (due to predictor compatibility issues), the Rescale implementation from Ancona et al. (2017) was used instead. The reference was set to the mean sequence pattern (PSSM) of the data set. We used either the absolute value of the saliency scores (‘DeepLIFT’) or the signed scores (‘DeepLIFT (Signed)’). Tested for prediction tasks: MNIST (Figure 2), APA (Figure 3), 5’ UTR (Figure 4) and PPI (Figure 5).

Code: (RevealCancel; Shrikumar et al., 2017) https://github.com/kundajelab/deeplift. (Rescale; Anconda et al., 2018) https://github.com/marcoancona/DeepExplain.

#### DeepSHAP

(Lundberg et al., 2017) SHAP DeepExplainer (‘DeepSHAP’) was executed for each input pattern with 100 reference pattern sampled uniformly from the data set. We used either the absolute value of the saliency scores (‘DeepSHAP’) or the signed scores (‘DeepSHAP (Signed)’). Tested for prediction tasks: MNIST (Figure 2), APA (Figure 3), 5’ UTR (Figure 4) and PPI (Figure 5).

Code: https://github.com/slundberg/shap.

#### TorchRay

(Fong et al., 2019) The extremal preservation/perturbation method from the TorchRay package was used in multiple benchmarks. We tested both the perturbation operator where masked pixels/letters are zeroed (‘TorchRay Fade’) and blurred by a gaussian (‘TorchRay Blur’). For the sequence-predictive tasks, we changed the code base to support 1-dimensional patterns. In all tasks, we set the blur operator *σ* to 3, the mask operator *σ* was set to 5 and the ‘step’ argument was set to 2. The ‘area’ was set to 0.1. Tested for prediction tasks: MNIST (Figure 2) and PPI (Figure 5).

Code: https://github.com/facebookresearch/TorchRay.

#### Saliency Model

(Dabkowski et al., 2017) The Saliency Model-method was used in multiple benchmarks. For the sequence-predictive tasks, we changed the code base to support 1-dimensional input patterns. The ‘encoder’-part of the network was not pre-trained and the ‘selector’-component was not used. In all tasks, the blur kernel size was set to 9 and *σ* was set to 3. For every training iteration, blur was used 50% of the time as the perturbation operator and fade (zeroing features) otherwise. Tested for prediction tasks: MNIST (Figure 2) and PPI (Figure 5).

Code: https://github.com/PiotrDabkowski/pytorch-saliency.

#### Sufficient Input Subsets

(Carter et al., 2020) Sufficient Input Subsets (SIS) was included in the benchmark of the protein binder prediction task. The SIS threshold was set to the 50th percentile of the predicted scores (𝒫 (***x***)) across the data set. For input patterns (dimer pairs) with non-masked predictions below this threshold, we used a dynamic threshold of 0.8 times the non-masked predicted score. We tested multiple masking schemes: (1) Replacing masked residue positions with the mean letter value (‘SIS (Mean)’) and (2) Replacing masked residue positions with letters sampled from the marginal letter distribution, repeating the sampling process *X* times, and calculating the mean prediction across all *X* samples (‘SIS (*X* Sample)’). Tested for prediction tasks: PPI (Figure 5).

Code: https://github.com/google-research/google-research/tree/master/sufficient_input_subsets.

#### Scrambler (Inclusion)

Scrambler network trained to minimize Equation 2 using the temperature-based perturbator of Equation 1, with background 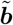 set to the mean input pattern across the training set. The network was trained between 20-50 epochs depending on task (when the validation error started to saturate).

#### Scrambler (Occlusion)

Scrambler network trained to minimize Equation 5 using the temperature-based perturbator of Equation 3, with background 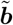 set to the mean input pattern across the training set.

#### Scrambler (Inclusion, Siamese)

Included in the PPI benchmark (Figure 5). Given pairs of input patterns ***x***_***1***_ and ***x***_***2***_, we optimize 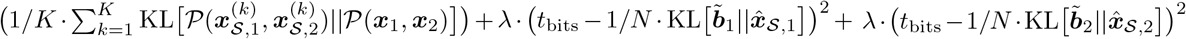, where 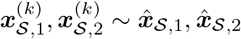 (letter-by-letter Gumbel-sampling) and 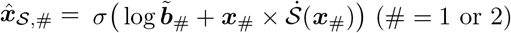. 𝒮 is a (siamese) Scrambler network.

#### Scrambler (Inclusion, Siamese, *X* Samples)

By default the Scrambler networks are trained by drawing 32 Gumbel samples from each scrambled PSSM in the training batch. In these versions, we instead draw *X* samples for each pattern in the training batch.

#### Scrambler (Occlusion, Joint)

Included in the PPI benchmark (Figure 5). Given pairs of input patterns ***x***_***1***_ and ***x***_***2***_, we optimize 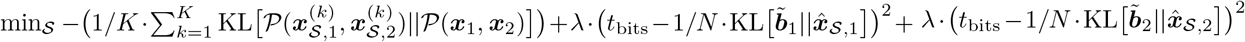, where 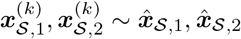 (letter-by-letter Gumbel-sampling) and 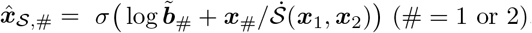. 𝒮 is a two-input Scrambler network.

#### Scrambler (Occlusion, Siamese)

Included in the PPI benchmark (Figure 5). Given pairs of input patterns ***x***_1_ and ***x***_2_, we optimize 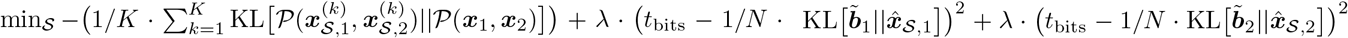, where 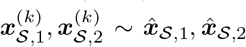 (letter-by-letter Gumbel-sampling) and 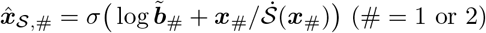. 𝒮 is a (siamese) Scrambler network.

#### Scrambler (Inclusion, Cont. Mean Bg)

Optimizes the same objective as the Inclusion-Scrambler, but uses a continuous interpolation perturbation with the mean training pattern 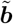 instead of the temperature-based perturbator: 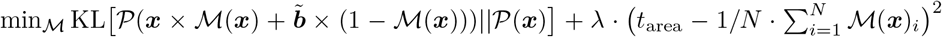, where ℳ (***x***) is a real-valued, sigmoid mask generator. Internally, network ℳ is identical to the Scrambler.

#### Scrambler (Inclusion, Cont. No Bg)

Optimizes the same objective as the Inclusion-Scrambler, but uses a continuous interpolation perturbation with the all-zero pattern instead of the temperature-based perturbator: 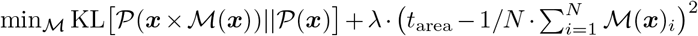, where ℳ (***x***) is a real-valued, sigmoid mask generator.

#### Scrambler (Inclusion, Cont. Rand Bg)

Optimizes the same objective as the Inclusion-Scrambler, but uses a continuous interpolation perturbation with a random letter pattern ℝ ∈ 0, 1^*N×M*^ instead of the temperature-based perturbator: 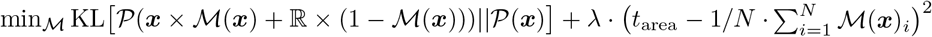, where ℳ (***x***) is a real-valued, sigmoid mask generator.

#### Scrambler (Occlusion, Cont. Mean Bg)

Optimizes the same objective as the Occlusion-Scrambler, but uses a continuous interpolation perturbation with the mean training pattern 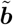 instead of the temperature-based perturbator: 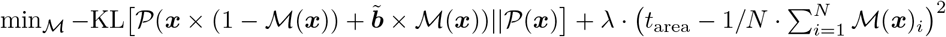, where ℳ (***x***) is a real-valued, sigmoid-restricted mask generator.

#### Scrambler (Inclusion, Per Example)

Optimizes the same objective as the Inclusion-Scrambler, but does not train a generative network to predict importance scores. Instead, this method optimizes a set of importance scores ***s*** for each pattern ***x*** that we wish to interpret individually: 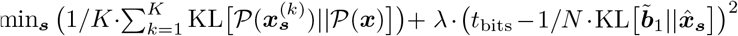, where 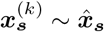 (letter-by-letter Gumbel-sampling) and 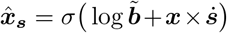. 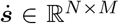 is a channel-broadcasted copy of the importance scores ***s*** ∈ ℝ^*N*^. We obtain ***s*** from a vector of real-valued (trainable) numbers that we instance-normalize and apply the softplus activation on. We train ***s*** until convergence using the Adam optimizer with learning rate = 0.01, *β*_1_ = 0.5 and *β*_2_ = 0.9.

#### Scrambler (Inclusion, Per Example, Cont. Mean Bg)

Optimizes the same objective as the Per-example Inclusion-Scrambler, but uses a continuous interpolation perturbation with the mean training pattern 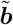 instead of the temperature-based perturbator: 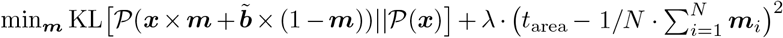, where ***m*** ∈ [0, 1]^*N*^ is a continuous sigmoid-restricted mask pattern.

#### Scrambler (Inclusion, Per Example, Cont. No Bg)

Optimizes the same objective as the Per-example Inclusion-Scrambler, but uses a continuous interpolation perturbation with the all-zero pattern instead of the temperature-based perturbator: 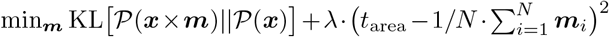, where ***m*** ∈ [0, 1]^*N*^ is a continuous sigmoid-restricted mask pattern.

#### Scrambler (Inclusion, Per Example, Cont. Rand Bg)

Optimizes the same objective as the Per-example Inclusion-Scrambler, but uses a continuous interpolation perturbation with a random letter pattern ℝ ∈ 0, 1^*N×M*^ instead of the temperature-based perturbator: 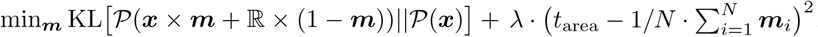, where ***m*** ∈ [0, 1]^*N*^ is a continuous sigmoid-restricted mask pattern.

#### Scrambler (Occlusion, Per Example, Cont. Mean Bg)

Optimizes the same objective as the Occlusion-Scrambler (for a single pattern), but uses a continuous interpolation perturbation with the mean training pattern 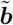 instead of the temperature-based perturbator: 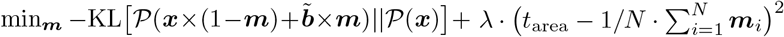, where ***m*** ∈ [0, 1]^*N*^ is a continuous sigmoid-restricted mask pattern.

#### Zero (Inclusion)

Optimizes the same high-level objective as the Scrambler, but this model uses a perturbation operator similar to that of L2X (Chen et al., 2018) and INVASE (Yoon et al., 2018) where de-selected features are zeroed. Specifically, Zero (Inclusion) optimizes 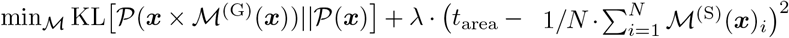, where ℳ is a binary 0*/*1 mask generator. The continuous-valued mask ℳ ^(S)^(***x***) and binarized mask ℳ ^(G)^(***x***) are obtained by applying sigmoid activations and Gumbel sampling on the output vector of ℳ respectively. Internally, the network ℳ is identical to the Scrambler.

#### Zero (Occlusion)

Optimizes 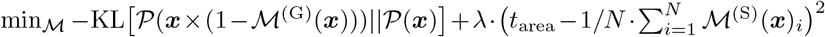, where ℳ is a binary 0*/*1 mask generator. See Zero (Inclusion) for more details.

#### Zero (Occlusion, Joint)

Included in the PPI benchmark (Figure 5). Given pairs of input patterns ***x***_1_ and ***x***_2_, we optimize 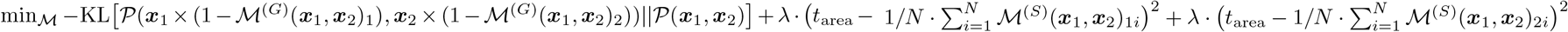, where ℳ is a pair-wise binary 0*/*1 mask generator. See Zero (Inclusion) for more details.

#### Zero (Inclusion, Siamese)

Included in the PPI benchmark (Figure 5). Given pairs of input patterns ***x***_1_ and ***x***_2_, we optimize 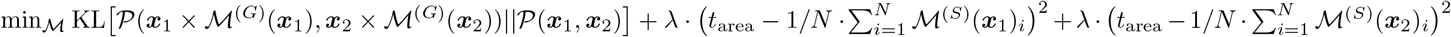, where ℳ is a binary 0*/*1 mask generator. See Zero (Inclusion) for more details.

#### Zero (Occlusion, Siamese)

Included in the PPI benchmark (Figure 5). Given pairs of input patterns ***x***_1_ and ***x***_2_, we optimize 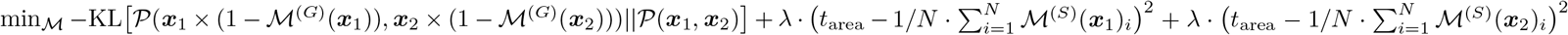, where ℳ is a binary 0*/*1 mask generator. See Zero (Inclusion) for more details.

#### Rand (Inclusion)

Optimizes the same high-level objective as the Scrambler, but this model uses a perturbation operator where de-selected features are replaced with a random nucleotide (or residue). Specifically, Rand (Inclusion) optimizes 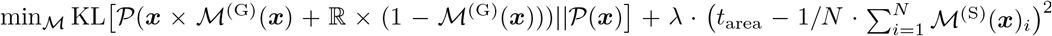, where ℳ is a binary 0*/*1 mask generator and ℝ is a random letter pattern. See Zero (Inclusion) for more details.

#### Rand (Occlusion)

Optimizes 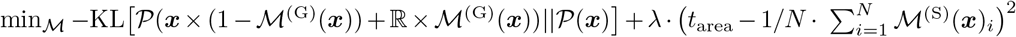, where ℳ is a binary 0*/*1 mask generator and ℝ is a random letter pattern. See Zero (Inclusion) for more details.

#### Rand (Occlusion, Joint)

Included in the PPI benchmark (Supplementary Fig. S5). Given pairs of input patterns ***x***_1_ and ***x***_2_, we optimize 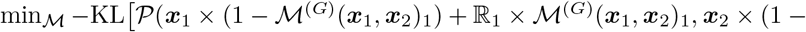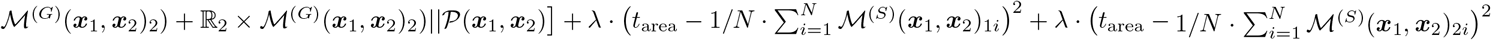, where ℳ is a pair-wise binary 0*/*1 mask generator and ℝ_1_, ℝ_2_ are random letter patterns. See Zero (Inclusion) for more details.

#### Rand (Inclusion, Siamese)

Included in the PPI benchmark (Supplementary Fig. S5). Given pairs of input patterns ***x***_1_ and ***x***_2_, we optimize 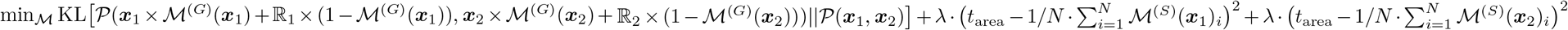, where ℳ is a binary 0*/*1 mask generator and ℝ_1_, ℝ_2_ are random letter patterns. See Zero (Inclusion) for more details.

#### Rand (Occlusion, Siamese)

Included in the PPI benchmark (Supplementary Fig. S5). Given pairs of input patterns ***x***_1_ and ***x***_2_, we optimize 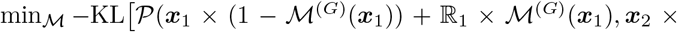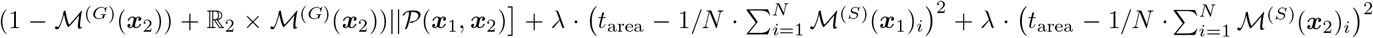, where ℳ is a binary 0*/*1 mask generator and ℝ_1_, ℝ_2_ are random letter patterns. See Zero (Inclusion) for more details.

### Benchmarking Details

#### Synthetic IF uORF 5’ UTR Datasets

Synthetic IF uORF 5’ UTR sequences were generated for feature attribution with varying numbers of IF uORFS by selecting sequences from the Optimus 5-Prime *egfp unmod 1* dataset (Sample et al., 2019), which contained no upstream starts or stops and had MRL values that fell between the 5th and 10th percentile of the dataset. This set of sequences (*n* = 537) had the original sequence nucleotides replaced by “ATG” and “TAG” uniformly at random at possible IF positions as needed to create the number of IF uORFs for the dataset, keeping the rest of the UTR fixed. Four synthetic 5’ UTR datasets of *n* = 512 sequences were generated: one IF uORF by inserting an “ATG” and followed by a “TAG”, two IF uORFs by inserting an “ATG” followed by two “TAG”, two IF uORFs by inserting two “ATG” followed by “TAG,” and four IF uORFs by inserting two “ATG” followed by two “TAG”.

#### Hydrogen Bond Network Detection

The Chen et al. (2019) dimers were designed to have a minimum of 4 residues involved, contacting all four helices, with all heavy-atom donors and acceptors in the network satisfied. However, in later design steps for the hetero-dimers, these networks were slightly disrupted making complete recovery difficult using the original parameters. As such, slightly relaxed criteria from the original design specifications were used to recover as many HBNet residues as possible from the test set designed structures. HBNet residues were recovered from test coiled-coil dimers using PyRosetta (Chaudhury, Lyskov, and Gray 2010) and the HBNetStapleInterface protocol, with the settings min network size=3, min helices contacted by network=3, hb threshold=-0.3, and find only native networks=True (Boyken et al., 2016). The ref2015 score function was used during HBNet mover setup (Alford et al., 2017).

#### Mean ΔΔG Difference Calculation and Alanine Scanning

Computational Alanine scanning was carried out for all residues in a hetero-dimer pair using PyRosetta (Chaudhury, Lyskov, and Gray 2010). Each position was mutated to Alanine and repacked with the neighboring residues prevented from design and repacking. The InterfaceAnalyzerMover was used with set pack input as False and set pack separated as True to calculate the mean mutation *δδG* in Rosetta Energy Units (REU) at each position. For each attribution method, the eight most important residues per dimer were selected, and the difference of the mean ΔΔ*G* of the top residues and the mean ΔΔ*G* of all other residues was computed.

#### Per-Residue Score Function Breakdown and Substitution Scoring Matrix

Lysozyme and Sensor Histidine Kinase structures were predicted with trRosetta (Yang et al., 2020). Each *de novo* structure was provided by (Anishchenko et al., 2020). All structures were relaxed using fast relax with PyRosetta (Chaudhury, Lyskov, and Gray 2010), scored using the *beta_nov16* score function (Alford et al., 2017), and broken down into individual residue contributions. This was done ten times per structure, and the average residue contributions were calculated.

## Availability of Data and Code

All code is available at http://www.github.com/johli/scrambler. External software and data used in this study are listed in the Methods section.

