## Supplemental Information for "Interpreting Neural Networks for Biological Sequences by Learning Stochastic Masks"

Table S1: An overview of interpretation methods based on masking. The columns describe (in order): Interpretation Algorithm – Whether the method uses a generative model to infer masks, or whether the masks are optimized individually; Masking Operator – Whether masked features are faded, blurred, zeroed or replaced with sampled values from either marginal distributions or from a generator ("counterfactual"); Application – What application the method was demonstrated on.

| | Interpretation Algorithm | Masking Operator | Application |
|---:|---|---|---|
| Scrambler Network (this paper) | Generative Model | Samples | Bio |
| Fong et al. (2017) (Fong et al., 2017) | Per-example SGD | Fade, Blur | Comp Vis |
| Dabkowski et al. (2017) (Dabowski et al., 2017) | Generative Model | Fade, Blur | Comp Vis |
| Yoon et al. (2018) (INVASE) (Yoon et al., 2017) | Generative Model | Zero | Comp Vis, NLP |
| Chen et al. (2018) (L2X)(Chen et al., 2018) | Generative Model | Zero | Comp Vis, NLP |
| Carter et al. (2019) (SIS) (Carter et al., 2019) | Per-example Recursion | Mean, Samples | Comp Vis, NLP, Bio |
| Zintgraf et al. (2017) (Zintgraf et al., 2017) | Per-example Sampling | Samples | Comp Vis |
| Chang et al. (2018) (FIDO-CA) (Chang et al., 2018) | Per-example SGD | Counterfactual | Comp Vis |

*Corresponding author

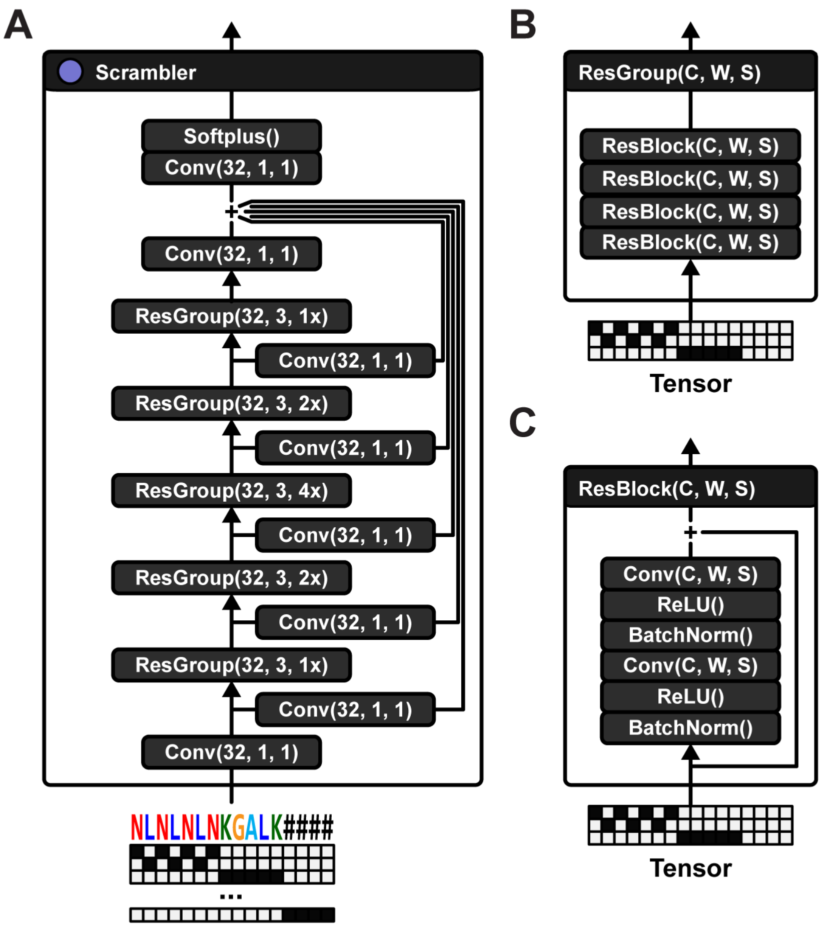


Figure S1: Scrambling neural networks. (A) Scrambler network architecture. The network is based on groups of residual blocks. This particular network configuration has 5 groups of residual blocks, with 32 channels, filter width 3 and varying dilation factor (1x, 2x, 4x, 2x, 1x). There is a skip connection (single convolutional layer with 32 channels and filter width 1) before each residual group. All skip connections are added together with the output of the final residual group. A softplus activation is applied to the final tensor in order to get importance scores that are strictly larger than 0. (B) Each residual group consists of 4 identical residual blocks connected in series. (C) Each residual block consists of 2 dilated convolutions, each preceded by batch normalization and ReLU activations. A skip connection adds the input tensor to the output of the final convolution.

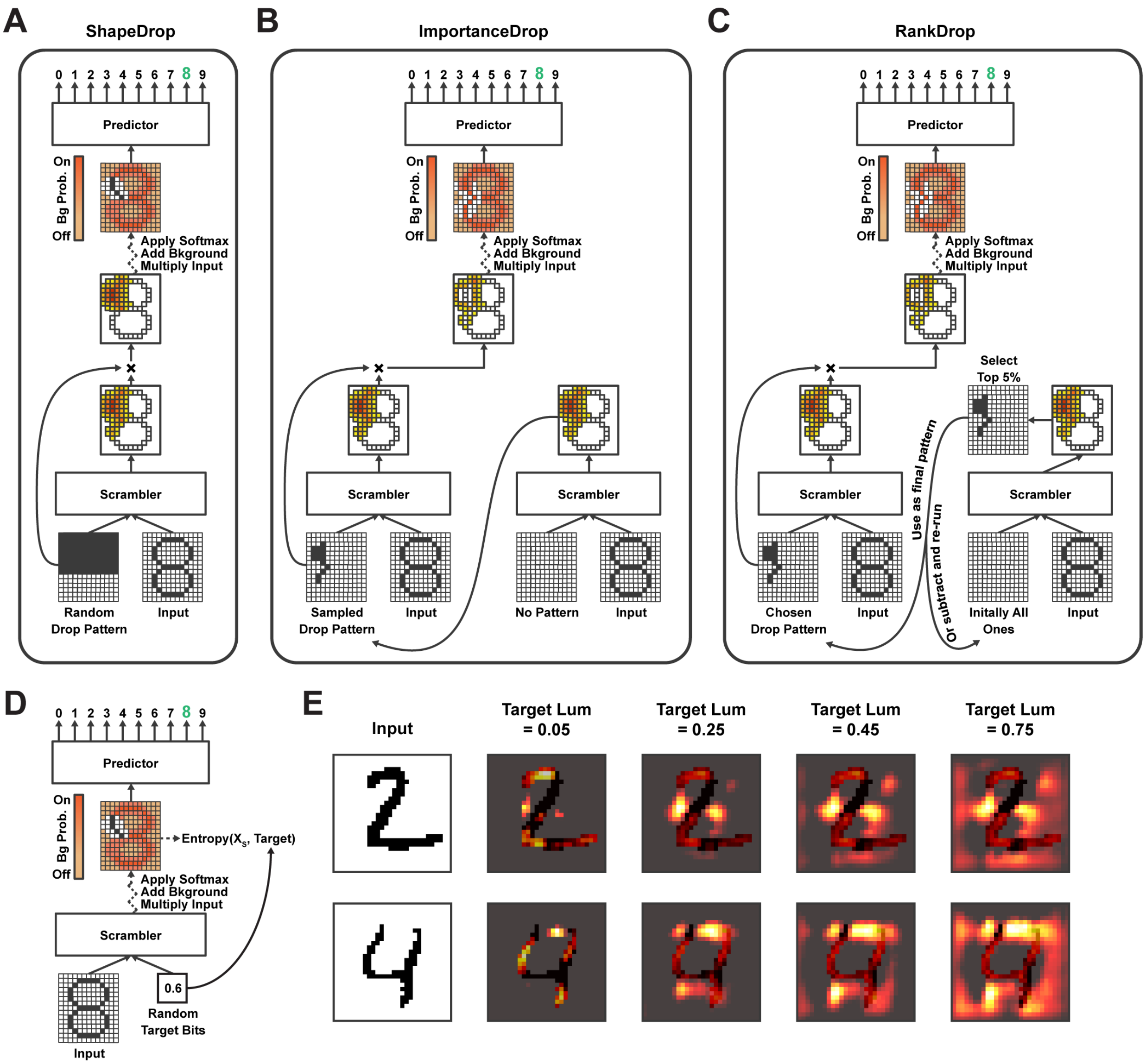


Figure S2: Scrambler extension MNIST demonstrations. (A) Uniformly random mask dropout training procedure, which teaches the Scrambler to find alternate salient feature sets. For each input pattern, we sample a random dropout pattern containing squares of varying width. The pattern is multiplied with the predicted importance scores, effectively zeroing out certain regions (forcing the background distribution to be used). The dropout pattern is also passed as additional input to the Scrambler (it is concatenated along the channel dimension), allowing the network to learn which other feature set to choose. (B) Biased dropout training procedure. Instead of randomly sampling dropout patterns, we first let the Scrambler predict importance scores with an all-ones dropout pattern (no dropout), which we use to form an importance sampling distribution. We then sample a dropout pattern and use it for training the same way we trained on uniformly random patterns. (C) Another biased dropout training approach. We first use the Scrambler to predict the importance scores given the all-ones dropout pattern as input (no dropout). The top 5% most important features are subtracted from the all-ones pattern. Then, with a certain probability, we either re-run the Scrambler on this updated pattern (repeating the previous steps), or we end the loop and choose this as our final dropout pattern to train on. (D) Procedure for training the Scrambler to dynamically change the entropy of its solutions. Instead of fitting the network to a constant KL-divergence of its scrambled input distribution, we here randomly sample bit values and use them both as input to the network and as the target for the conservation penalty. The bit value is broadcasted and concatenated along the input channel dimension. (E) Example attributions of MNIST digit '2' and '4' with dynamically resized feature sets, by passing increasingly large target 'lum' values as input to the Scrambler ('lum' values are normalized KL-divergence bits, see Methods).

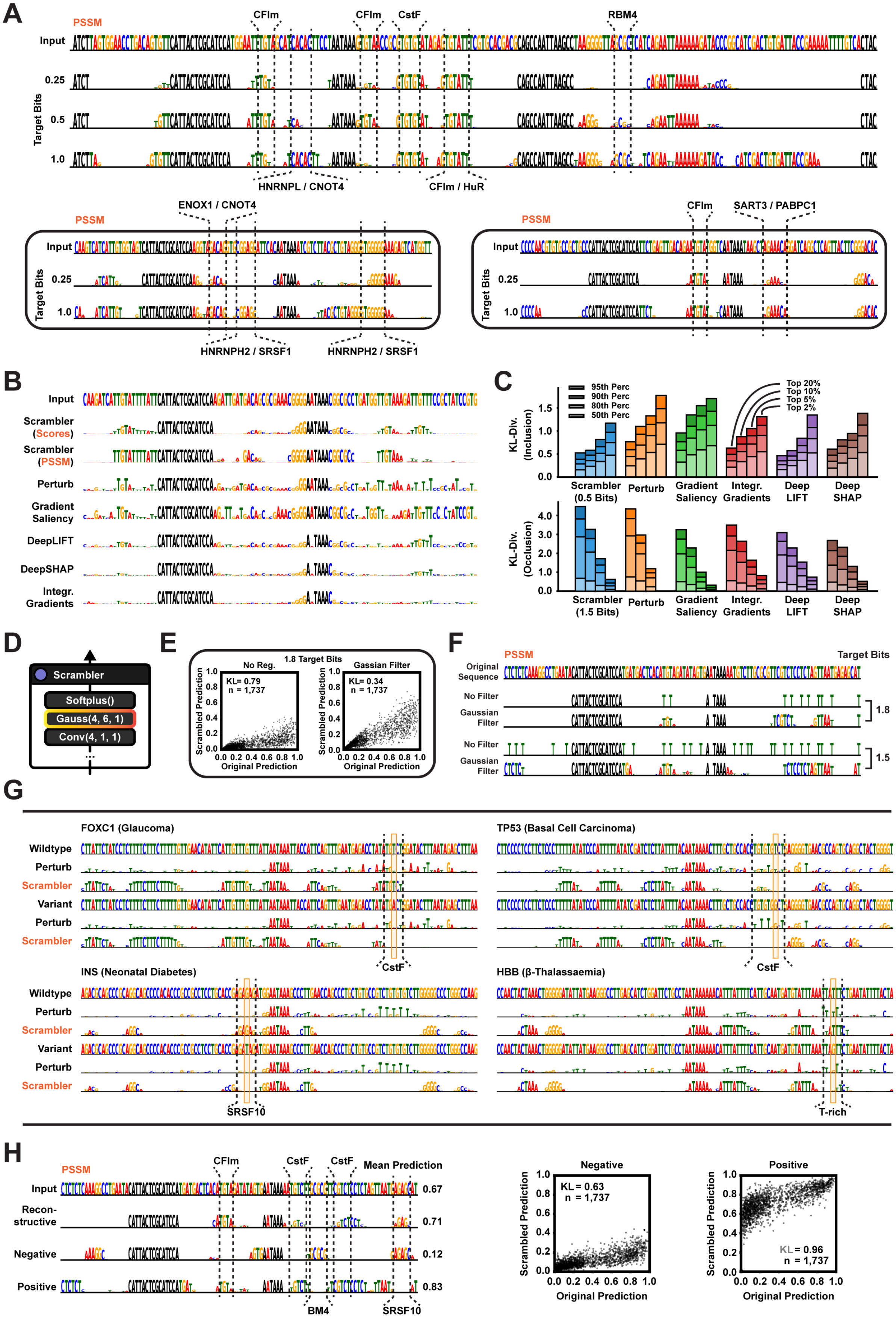
A
PSSM
Input
Target Bits
0.25
0.5
1.0
CFIm
CFIm
CstF
RBM4
HNRNPL / CNOT4
CFIm / HuR
ENOX1 / CNOT4
HNRNPH2 / SRSF1
HNRNPH2 / SRSF1
SART3 / PABPC1
B
Input
Scrambler (Scores)
Scrambler (PSSM)
Perturb
Gradient Saliency
DeepLIFT
DeepSHAP
Integr. Gradients
C
95th Perc
90th Perc
80th Perc
50th Perc
Top 20%
Top 10%
Top 5%
Top 2%
KL-Div. (Inclusion)
KL-Div. (Occlusion)
Scrambler (0.5 Bits)
Scrambler (1.5 Bits)
Perturb
Gradient Saliency
Integr. Gradients
Deep LIFT
Deep SHAP
D
Scrambler
Softplus()
Gauss(4, 6, 1)
Conv(4, 1, 1)
E
1.8 Target Bits
No Reg.
Gassian Filter
KL= 0.79
n = 1,737
KL= 0.34
n = 1,737
Scrambled Prediction
Original Prediction
F
PSSM
Target Bits
Original Sequence
No Filter
Gaussian Filter
1.8
1.5
G
FOXC1 (Glaucoma)
TP53 (Basal Cell Carcinoma)
INS (Neonatal Diabetes)
HBB (β-Thalassaemia)
Wildtype
Perturb
Scrambler
Variant
CstF
SRSF10
T-rich
H
PSSM
Input
Recon-structive
Negative
Positive
Mean Prediction
0.67
0.71
0.12
0.83
CFIm
CstF
CstF
BM4
SRSF10
Negative
Positive
KL = 0.63
n = 1,737
KL = 0.96
n = 1,737
Scrambled Prediction
Original Prediction

Figure S3: APARENT attributions. (A) Example attribution of a strong polyadenylation signal sequence from the APARENT (Bogard et al., 2018) test set, using Inclusion-Scramblers trained with increasingly large $t_{\text{bits}}$ of conservation. Known regulatory motifs annotated. (B) Example attributions of a polyadenylation signal sequence, comparing different methods. (C) Inclusion and occlusion benchmark, comparing the Scrambler to Perturbation, Gradient Saliency (Simonyan et al., 2013), Integrated Gradients (Sundararajan et al., 2017), DeepLIFT (Rescale) (Shrikumar et al., 2017) and DeepSHAP (Lundberg et al., 2017) on the APARENT test set. In the inclusion benchmark (left), the top 20%, 10%, 5% and 2% most important nucleotide positions per sequence is kept fixed (according to each method) while other nucleotides are replaced with random samples. In the occlusion benchmark, these highest-scoring nucleotides are instead replaced with random samples while the remaining sequence is kept fixed. In each, we measure how much the prediction KL-divergence is either preserved or perturbed compared to the original sequences. (D) Non-trainable convolutional filter with a 1D gaussian kernel (filter width 6) is prepended to the final softplus activation function of the Scrambler. (E) APARENT isoform predictions of original sequences and of corresponding sampled sequences from the PSSMs generated by an Occlusion-Scrambler trained with $t_{\text{bits}} = 1.8$, with and without the Gaussian filter. (F) Example attributions of $t_{\text{bits}} = 1.8$- and 1.5 Occlusion-Scramblers, with and without the Gaussian filter. (G) Attributions of four human polyadenylation signals which are associated with known deleterious variants, comparing the Perturbation method to the reconstructive Inclusion-Scrambler ($t_{\text{bits}} = 0.25$ target bits) on hypothetical variants which have not been found in the population. Gene names and clinical condition associated with the PAS annotated above each sequence (Stacey et al., 2011; Medina-Trillo et al., 2016; Altay et al., 1991; Garin et al., 2010). In each of the four examples, the Scrambler correctly detects the loss of the presumed RBP binding site or otherwise important motif due to each respective variant (loss of the CstF binding motif in *FOXC1*, *TP53*; loss of the SRSF10 binding motif in *INS*; loss of the T-rich DSE motif in *HBB*). (H) Left: Example attributions of a medium-strength polyadenylation signal sequence, using three Scramblers which have been optimized for different objectives: (Reconstructive features) reconstructing the original prediction, (Negative features) minimizing the prediction, and (Positive features) maximizing the prediction. Right: APARENT isoform predictions of original sequences and of corresponding sampled sequences from the PSSMs generated by the Negative-feature and Positive-feature Scrambler respectively.

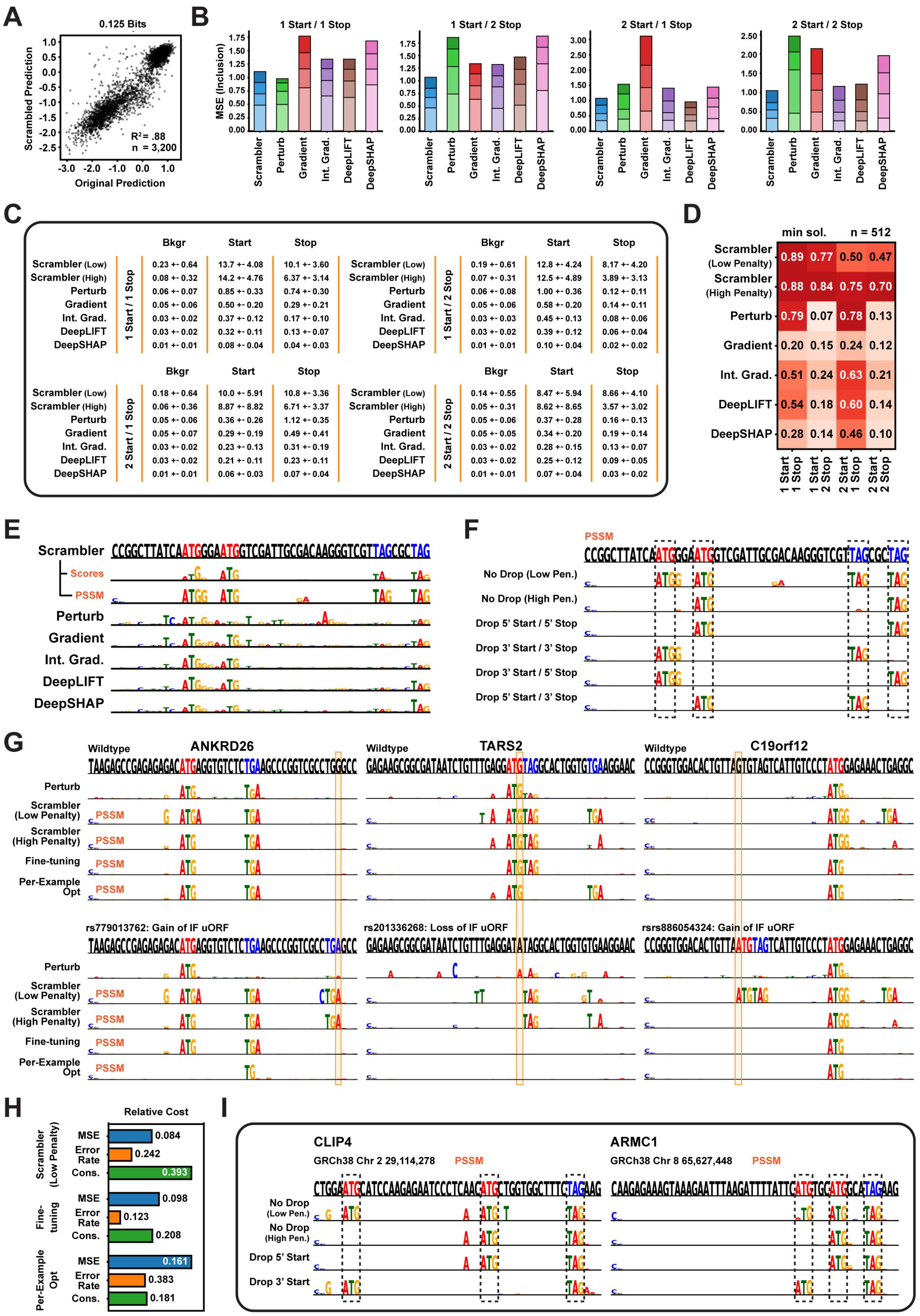


| 1 Start / 1 Stop | Bkgr | Start | Stop |
|---|---|---|---|
| Scrambler (Low) | 0.23 +- 0.64 | 13.7 +- 4.08 | 10.1 +- 3.60 |
| Scrambler (High) | 0.08 +- 0.32 | 14.2 +- 4.76 | 6.37 +- 3.14 |
| Perturb | 0.06 +- 0.07 | 0.85 +- 0.33 | 0.74 +- 0.30 |
| Gradient | 0.05 +- 0.06 | 0.50 +- 0.20 | 0.29 +- 0.21 |
| Int. Grad. | 0.03 +- 0.02 | 0.37 +- 0.12 | 0.17 +- 0.10 |
| DeepLIFT | 0.03 +- 0.02 | 0.32 +- 0.11 | 0.13 +- 0.07 |
| DeepSHAP | 0.01 +- 0.01 | 0.08 +- 0.04 | 0.04 +- 0.03 |

| 1 Start / 2 Stop | Bkgr | Start | Stop |
|---|---|---|---|
| Scrambler (Low) | 0.19 +- 0.61 | 12.8 +- 4.24 | 8.17 +- 4.20 |
| Scrambler (High) | 0.07 +- 0.31 | 12.5 +- 4.89 | 3.89 +- 3.13 |
| Perturb | 0.06 +- 0.08 | 1.00 +- 0.36 | 0.12 +- 0.11 |
| Gradient | 0.05 +- 0.06 | 0.58 +- 0.20 | 0.14 +- 0.11 |
| Int. Grad. | 0.03 +- 0.03 | 0.45 +- 0.13 | 0.08 +- 0.06 |
| DeepLIFT | 0.03 +- 0.02 | 0.39 +- 0.12 | 0.06 +- 0.04 |
| DeepSHAP | 0.01 +- 0.01 | 0.10 +- 0.04 | 0.02 +- 0.02 |

| 2 Start / 1 Stop | Bkgr | Start | Stop |
|---|---|---|---|
| Scrambler (Low) | 0.18 +- 0.64 | 10.0 +- 5.91 | 10.8 +- 3.36 |
| Scrambler (High) | 0.06 +- 0.36 | 8.87 +- 8.82 | 6.71 +- 3.37 |
| Perturb | 0.05 +- 0.06 | 0.36 +- 0.26 | 1.12 +- 0.35 |
| Gradient | 0.05 +- 0.07 | 0.29 +- 0.19 | 0.49 +- 0.41 |
| Int. Grad. | 0.03 +- 0.02 | 0.23 +- 0.13 | 0.31 +- 0.19 |
| DeepLIFT | 0.03 +- 0.02 | 0.21 +- 0.11 | 0.23 +- 0.11 |
| DeepSHAP | 0.01 +- 0.01 | 0.06 +- 0.03 | 0.07 +- 0.04 |

| 2 Start / 2 Stop | Bkgr | Start | Stop |
|---|---|---|---|
| Scrambler (Low) | 0.14 +- 0.55 | 8.47 +- 5.94 | 8.66 +- 4.10 |
| Scrambler (High) | 0.05 +- 0.31 | 8.62 +- 8.65 | 3.57 +- 3.02 |
| Perturb | 0.05 +- 0.06 | 0.37 +- 0.28 | 0.16 +- 0.13 |
| Gradient | 0.05 +- 0.05 | 0.34 +- 0.20 | 0.19 +- 0.14 |
| Int. Grad. | 0.03 +- 0.02 | 0.28 +- 0.15 | 0.13 +- 0.07 |
| DeepLIFT | 0.03 +- 0.02 | 0.25 +- 0.12 | 0.09 +- 0.05 |
| DeepSHAP | 0.01 +- 0.01 | 0.07 +- 0.04 | 0.03 +- 0.02 |

Figure S4: Optimus 5-Prime attributions. (A) Optimus 5-Prime (Sample et al., 2019) MRL predictions of test sequences compared to predictions of sequence samples drawn the corresponding Scrambler PSSMs ($t_{\text{bits}} = 0.125$). (B) Inclusion benchmark on synthetic test sequences. For 1 start / 1 stop, 6 nucleotides are fixed, for 1 start / 2 stop and 2 start / 1 stop, 9 nucleotides are fixed and for 2 start / 2 stop, the 12 most important nucleotides are fixed. Remaining nucleotides are replaced with random samples and the prediction KL-divergence is compared to the original sequences is used to evaluate reconstruction quality. (C) Mean importance scores (and standard deviation) for positions containing either an arbitrary nucleotide ('Bkgr'), a start codon or a stop codon, measured across the synthetic test of sequences per logic function. (D) Average recall for finding one of the start codons and one of the stop codons in the 6 most important nucleotides, as identified by each method, measured across the synthetic test sets. (E) Example attribution of a 5' UTR sequence with two IF start codons (marked with red letters in the top sequence) and two IF stop codons (marked with blue letters). (F) Example attributions using a Scrambler network trained with the mask dropout procedure (see Methods for details). By masking different parts of the sequence, the Scrambler discovers other valid salient feature sets (IF uORFs). (G) Left: Attribution of a ClinVar variant, *rs779013762*, in the *ANKRD26* 5' UTR, which is predicted by Optimus 5-Prime to be a functionally silent mutation. The variant creates an IF uORF overlapping an existing IF uORF. The per-example fine-tuning step (which starts from the Low entropy penalty-Scrambler scores) finds a minimal salient feature set in the variant sequence (one IF uORF), while the per-example optimization (which starts from randomly initialized scores) gets stuck in a local minimum. Middle: Attribution of a ClinVar variant, *rs201336268*, in the *TARS2* 5' UTR, which destroys two overlapping IF uORFs and is predicted to lead to upregulation. Both the fine-tuning step and the independent per-example optimization finds that no features are important in the variant sequence (both IF uORFs were removed by the variant and a fully random sequence has on average the same predicted MRL as the variant sequence). The Perturbation method has trouble explaining either of these variants due to saturation effects of the multiple IF stop codons. Right: Attribution of a rare variant, *rs886054324*, in the *C19orf12* 5' UTR, which creates two IF uORFs overlapping a strong OOF uAUG (hence a silent mutation). All attribution methods identify the OOF uAUG as the major determinant, however the Low entropy penalty-Scrambler incorrectly marks an (unmatched) stop codon in the wildtype sequence as important. Both the High entropy penalty-Scrambler and the fine-tuning step based off the Low penalty-Scrambler correctly filters the stop codon. (H) Benchmarking results on the 1 Start / 2 Stop dataset (see Supplementary Fig. S4B-D), comparing the Low entropy penalty-Scrambler network to running per-example fine-tuning of those scores and to the baseline method of optimizing each example from randomly initialized scores. Reported are the mean squared error between predictions on original and scrambled sequences ('MSE'), the error rate ($1 - \textit{Accuracy}$) of not finding one Start codon and one Stop codon in the top 6 nt ('Error Rate'), and the mean per-nucleotide KL-divergence between the scrambled PSSM and the background PSSM ('Conservation'). (I) Example Scrambler attributions with the mask dropout mechanism on two native human 5' UTRs.

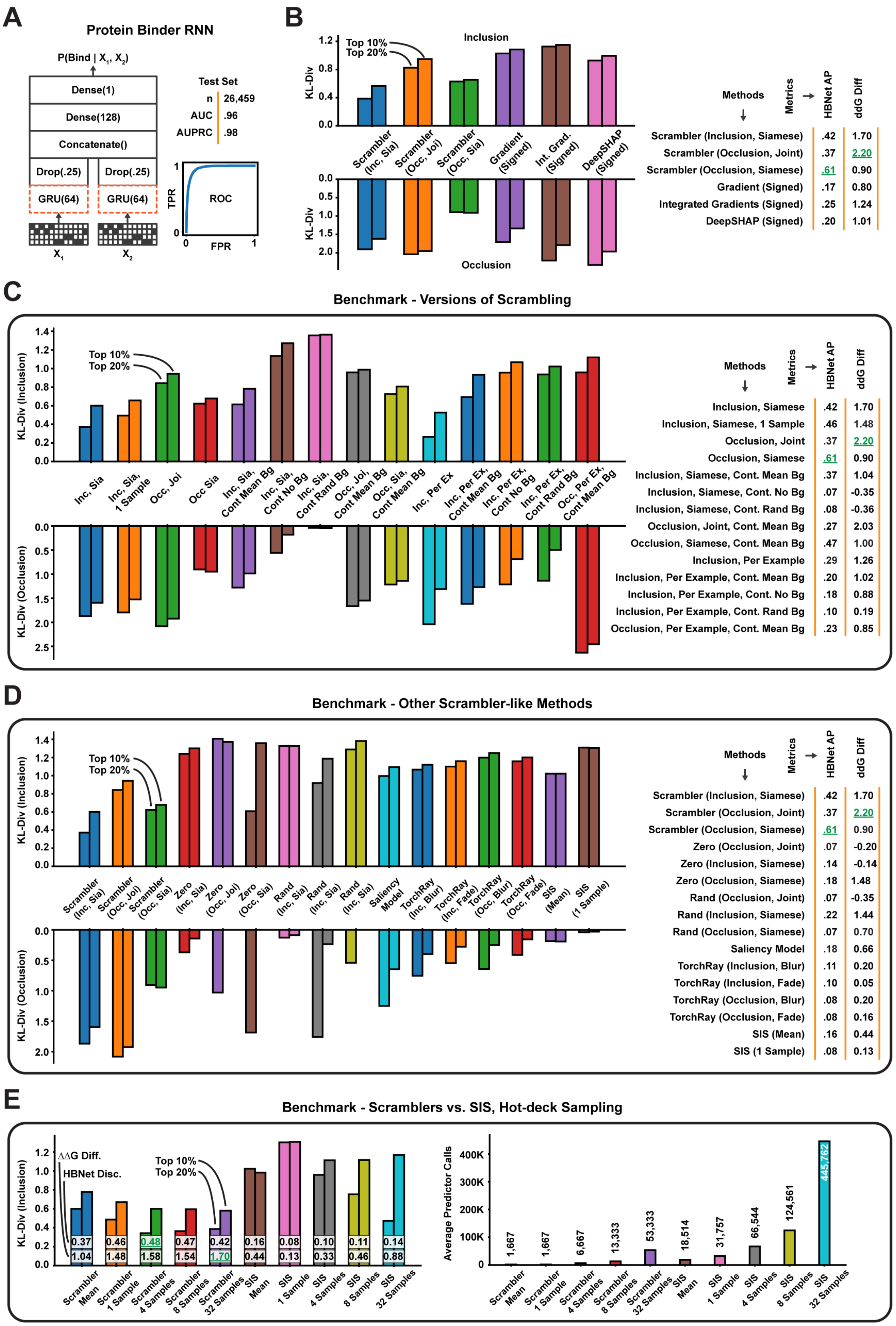

A
Protein Binder RNN
P(Bind | X1, X2)
Dense(1)
Dense(128)
Concatenate()
Drop(.25)
Drop(.25)
GRU(64)
GRU(64)
X1
X2
Test Set
n 26,459
AUC .96
AUPRC .98
TPR
FPR
ROC
B
Inclusion
Occlusion
KL-Div
Top 10%
Top 20%
Scrambler (Inc, Sia)
Scrambler (Occ, Joi)
Scrambler (Occ, Sia)
Gradient (Signed)
Int. Grad. (Signed)
DeepSHAP (Signed)
Methods
Metrics
HBNet AP
ddG Diff
Scrambler (Inclusion, Siamese) .42 1.70
Scrambler (Occlusion, Joint) .37 2.20
Scrambler (Occlusion, Siamese) .61 0.90
Gradient (Signed) .17 0.80
Integrated Gradients (Signed) .25 1.24
DeepSHAP (Signed) .20 1.01
C
Benchmark - Versions of Scrambling
KL-Div (Inclusion)
KL-Div (Occlusion)
Inc, Sia
Inc, Sia, 1 Sample
Occ, Joi
Occ Sia
Inc, Sia, Cont Mean Bg
Inc, Sia, Cont No Bg
Inc, Sia, Cont Rand Bg
Occ, Joi, Cont Mean Bg
Occ, Sia, Cont Mean Bg
Inc, Per Ex
Inc, Per Ex, Cont Mean Bg
Inc, Per Ex, Cont No Bg
Inc, Per Ex, Cont Rand Bg
Occ, Per Ex, Cont Mean Bg
Methods
Metrics
HBNet AP
ddG Diff
Inclusion, Siamese .42 1.70
Inclusion, Siamese, 1 Sample .46 1.48
Occlusion, Joint .37 2.20
Occlusion, Siamese .61 0.90
Inclusion, Siamese, Cont. Mean Bg .37 1.04
Inclusion, Siamese, Cont. No Bg .07 -0.35
Inclusion, Siamese, Cont. Rand Bg .08 -0.36
Occlusion, Joint, Cont. Mean Bg .27 2.03
Occlusion, Siamese, Cont. Mean Bg .47 1.00
Inclusion, Per Example .29 1.26
Inclusion, Per Example, Cont. Mean Bg .20 1.02
Inclusion, Per Example, Cont. No Bg .18 0.88
Inclusion, Per Example, Cont. Rand Bg .10 0.19
Occlusion, Per Example, Cont. Mean Bg .23 0.85
D
Benchmark - Other Scrambler-like Methods
Scrambler (Inc, Sia)
Scrambler (Occ, Joi)
Scrambler (Occ, Sia)
Zero (Inc, Sia)
Zero (Occ, Joi)
Zero (Occ, Sia)
Rand (Inc, Sia)
Rand (Inc, Sia)
Rand (Inc, Sia)
Saliency Model
TorchRay (Inc, Blur)
TorchRay (Inc, Fade)
TorchRay (Occ, Blur)
TorchRay (Occ, Fade)
SIS (Mean)
SIS (1 Sample)
Methods
Metrics
HBNet AP
ddG Diff
Scrambler (Inclusion, Siamese) .42 1.70
Scrambler (Occlusion, Joint) .37 2.20
Scrambler (Occlusion, Siamese) .61 0.90
Zero (Occlusion, Joint) .07 -0.20
Zero (Inclusion, Siamese) .14 -0.14
Zero (Occlusion, Siamese) .18 1.48
Rand (Occlusion, Joint) .07 -0.35
Rand (Inclusion, Siamese) .22 1.44
Rand (Occlusion, Siamese) .07 0.70
Saliency Model .18 0.66
TorchRay (Inclusion, Blur) .11 0.20
TorchRay (Inclusion, Fade) .10 0.05
TorchRay (Occlusion, Blur) .08 0.20
TorchRay (Occlusion, Fade) .08 0.16
SIS (Mean) .16 0.44
SIS (1 Sample) .08 0.13
E
Benchmark - Scramblers vs. SIS, Hot-deck Sampling
ΔΔG Diff.
HBNet Disc.
Top 10%
Top 20%
0.37 1.04
0.46 1.48
0.48 1.58
0.47 1.54
0.42 1.70
0.16 0.44
0.08 0.13
0.10 0.33
0.11 0.46
0.14 0.88
Scrambler Mean
Scrambler 1 Sample
Scrambler 4 Samples
Scrambler 8 Samples
Scrambler 32 Samples
SIS Mean
SIS 1 Sample
SIS 4 Samples
SIS 8 Samples
SIS 32 Samples
Average Predictor Calls
1,667
1,667
6,667
13,333
53,333
18,514
31,757
66,544
124,561
445,762

Figure S5: Heterodimer predictor attributions. (A) Protein heterodimer binder RNN predictor, which was trained on computationally designed (dimerizing) pairs for positive data and randomly paired binders as negative data (see Methods for details). The RNN consists of a shared GRU layer, a dropout layer, and two fully-connected layers applied to the concatenated GRU output vectors. The final output (sigmoid activation) is treated as the Bind / No Bind classification probability. (B) Supplemental benchmark of Gradient Saliency (Simonyan et al., 2013), DeepLIFT (Shrikumar et al., 2017) and DeepSHAP (Lundberg et al., 2017), using only the positive-valued importance scores. Left: Prediction KL-divergence of scrambled sequences compared to original test set sequences when either replacing all but the top X% most important amino acid residues with random samples (inclusion) or, conversely, when replacing the top X% nucleotides with random samples and keeping the remaining sequence fixed (occlusion). Right: Mean ddG Difference for the top 8 most important residues according to each method, measured across the test set, and HBNet (Maguire et al., 2018) Average Precision based on each method's importance scores. (C) Supplemental comparison of different versions of the Scrambling Neural Network (see Methods for a full description of each version). Left: KL-divergence benchmark based on the predictor RNN. Right: Mean ddG Differences and HBNet Discovery Precisions. (D) Supplemental comparison of other methods that optimize similar objectives as the Scrambler (see Methods for a full description of each method). Left: KL-divergence benchmark based on the predictor RNN. Right: Mean ddG Differences and HBNet Discovery Precisions. (E) Supplemental comparison between Scrambling Neural Networks and Sufficient Input Subsets (SIS) with 'hot-deck' sampled masking (the number of samples used at each iteration is varied from 1 to 32; see Methods for details). Left: KL-divergence benchmark based on the predictor RNN, Mean ddG Differences and HBNet Discovery Precisions annotated on top of the bar chart. Right: Average number of predictor queries used to interpret a single input pattern (for the Scrambler, this is the amortized cost of training divided by the number of test patterns interpreted).

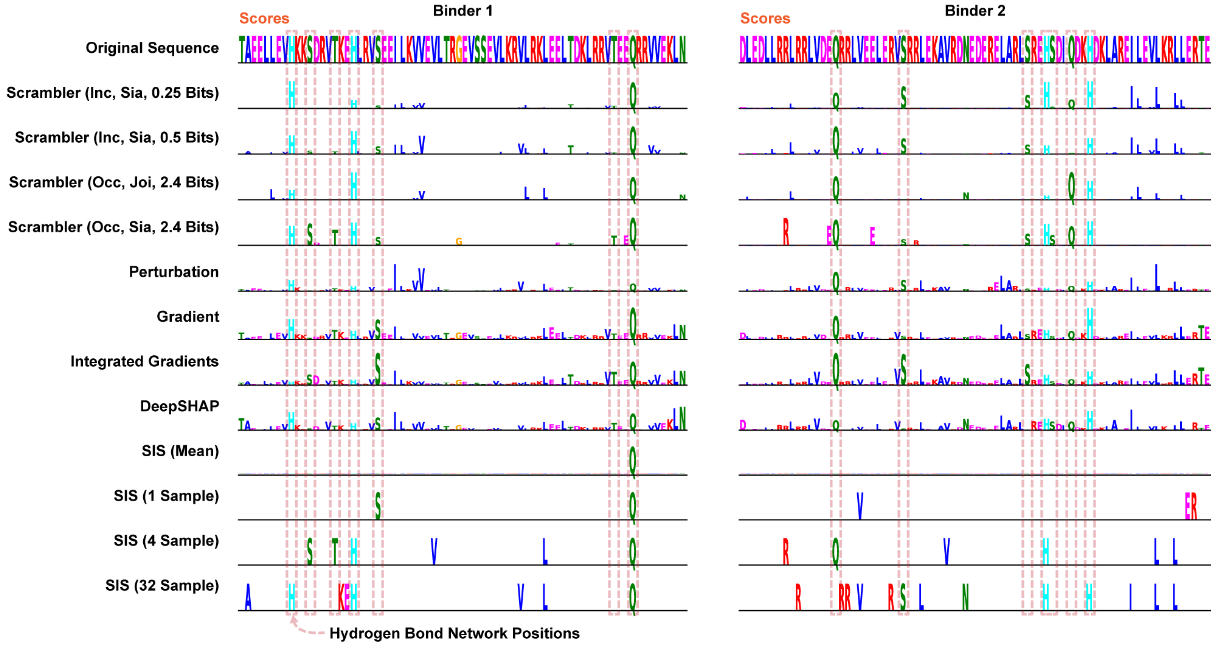


Figure S6: Heterodimer comparative attribution visualization. Example attributions of a designed heterodimer binder pair, for a selection of benchmarked methods.

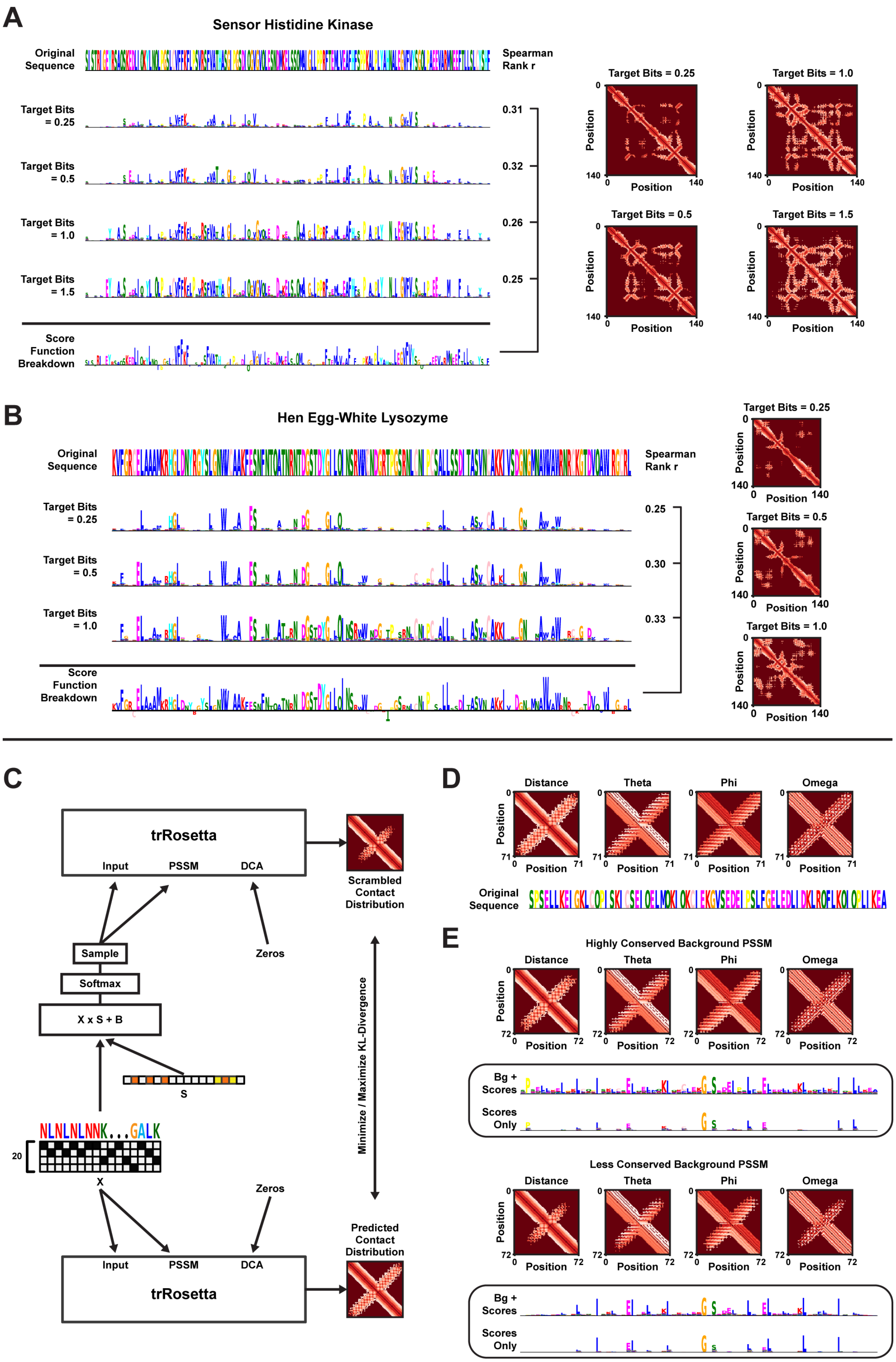
A
Sensor Histidine Kinase
Original Sequence
Spearman Rank r
Target Bits = 0.25
Target Bits = 0.5
Target Bits = 1.0
Target Bits = 1.5
0.31
0.32
0.26
0.25
Score Function Breakdown
Target Bits = 0.25
Target Bits = 1.0
Target Bits = 0.5
Target Bits = 1.5
Position
B
Hen Egg-White Lysozyme
Original Sequence
Spearman Rank r
Target Bits = 0.25
Target Bits = 0.5
Target Bits = 1.0
0.25
0.30
0.33
Score Function Breakdown
Position
C
trRosetta
Input
PSSM
DCA
Scrambled Contact Distribution
Sample
Softmax
X x S + B
Zeros
S
Minimize / Maximize KL-Divergence
NLNLNLNNK...GALK
20
X
Predicted Contact Distribution
D
Distance
Theta
Phi
Omega
Position
Original Sequence
E
Highly Conserved Background PSSM
Bg + Scores
Scores Only
Less Conserved Background PSSM

Figure S7: Native and *de novo* protein trRosetta attributions. (A) Four different Inclusion-PSSMs optimized to reconstruct the structural trRosetta (Yang et al., 2020) prediction of a Sensor Histidine Kinase. Each PSSM is optimized for increasingly larger $t_{\text{bits}}$. The bottom sequence logo represents the Rosetta score function breakdown per residue (−REU). Spearman r ranged between 0.25 and 0.32 when comparing the absolute numbers of Rosetta energy values to the optimized importance scores. Shown is also the average structure prediction for 512 samples. (B) Inclusion-Scrambled PSSMs of the Hen Egg-white Lysozyme. The PSSM was re-optimized for three different target conservation bits. Spearman r ranged between 0.25 and 0.33 compared to the Rosetta score function. (C) Architecture for per-example scrambling of a single protein sequence according to the contact distributions predicted by trRosetta. Here, we do not use a Multiple Sequence Alignment (MSA), but instead pass the Gumbel-sampled sequence to the PSSM input and an all-zeros matrix to the DCA input. Total KL-divergence between trRosetta-predicted distributions (distance and angle-grams) of the original sequence and samples drawn from the scrambled PSSM is either minimized or maximized (inclusion or occlusion respectively). (D) Reference sequence and predicted contact distribution for a hairpin protein engineered by Activation Maximization. (E) Top: Inclusion-PSSM of the engineered hairpin protein, obtained after optimization with a highly conserved background distribution (generated by HHblits (Remmert et al., 2020)). Bottom: Inclusion-PSSM of the engineered hairpin protein with a less conserved background distribution (smoothed with pseudo counts).
